# YAP1 defines an emergent, plastic population of relapsed small cell lung cancer

**DOI:** 10.1101/2025.10.21.683746

**Authors:** C. Allison Stewart, Kavya Ramkumar, Runsheng Wang, Yuanxin Xi, Alexa Halliday, Lixia Diao, Qi Wang, Alejandra Serrano, Sarah Groves, Simon Heeke, Azusa Tanimoto, Laura Kaiser, Whitney Lewis, Mukulika Bose, Pedro Da Rocha, Loukia Karacosta, Vito Quaranta, Jing Wang, Julie George, Luisa Maren Solis Soto, Bingnan Zhang, John V. Heymach, Lauren A. Byers, Carl M. Gay

## Abstract

Small cell lung cancer (SCLC) is an aggressive neuroendocrine malignancy characterized by rapid onset of chemoresistance and poor clinical outcomes. Transcriptional heterogeneity among treatment-naïve SCLC tumors underlies four transcriptional subtypes, each with distinct clinical vulnerabilities. Though previously hypothesized to delineate a distinct subtype, expression of YAP1 is largely absent from treatment-naïve, pure SCLC. To characterize relapsed SCLC, circulating tumor DNA, circulating tumor cells, and core needle biopsies from SCLC patients and preclinical models following resistance to standard-of-care therapies were analyzed. In contrast to treatment-naïve SCLC, these analyses reveal an emergent YAP1-positive cell population that coincides with treatment resistance. These YAP1-positive cells exhibit characteristics of drug tolerant persister cells, including senescence, stemness, and plasticity, as YAP1 positive cells largely abandon features characteristic of SCLC to adopt those of large-cell neuroendocrine carcinoma (LCNEC). As a result of this SCLC-like to LCNEC-like evolution, *YAP1*-positive cells lack several clinically relevant SCLC surface targets (i.e., DLL3, SEZ6), but are enriched for others (i.e., B7-H3, TROP2). We propose a model where YAP1 expressing cells emerge with SCLC treatment resistance and characterize a tenacious subpopulation capable of diverging from the treatment naïve lineage and adopting features to evade therapeutic response.

## Background

Small cell lung cancer (SCLC) is a high-grade neuroendocrine carcinoma that accounts for approximately 15% of all lung cancers diagnosed worldwide. While initially sensitive to platinum-based chemotherapy, SCLC is noteworthy for its resilience and resistance to therapy is virtually inevitable. As a result, despite initial chemotherapy response rates in excess of 70%, median survival for extensive-stage (ES)-SCLC patients is just over one year^1-3^. Further limiting progress in SCLC is that, in contrast to non-small cell lung cancer (NSCLC), patients are still subject to one-size-fits-all approaches with no standard predictive biomarkers.

SCLC molecular subtyping provides a framework to delineate inter-tumoral heterogeneity and, potentially, to personalize therapies based on subtype-specific vulnerabilities. Transcriptional analyses have revealed four major molecular subtypes – three defined by predominance of the transcription factors ASCL1 (SCLC-A), NEUROD1 (SCLC-N), and POU2F3 (SCLC-P), along with a fourth defined by inflammatory features (SCLC-I) such as antigen presenting machinery, T-cell infiltrate, and immune checkpoints^4^.

These subtypes differ not only in their tumor immune microenvironment (TIME), but other spectra including epithelial/mesenchymal and neuroendocrine/non-neuroendocrine features. As for the potential clinical relevance of these subtypes, retrospective analyses of Phase 3 clinical trials demonstrate that SCLC-I predicts those patients most likely to experience long-term benefit from immunotherapy^4-6^, while preclinical data point to unique vulnerabilities among each of the other three subtypes.

Several questions remain regarding the SCLC-derived transcriptional subtypes, however. One recurrent point of controversy has been the role of the transcriptional co-activator YAP1^7^. YAP1 and the related transcription factor TAZ have been well-established in other solid tumors as key factors in tumor initiation, treatment resistance, metastasis, and stemness^8^. Earlier attempts to classify SCLC into transcriptional subtypes suggested, based on primarily SCLC cell line models, that a rare, but distinct SCLC population was defined by the presence of YAP1--mutually exclusive with ASCL1/NEUROD1/POU2F3^9^. While YAP1 expression defines a subtype in other treatment-naive neuroendocrine neoplasms (e.g., pulmonary large cell neuroendocrine carcinoma [LCNEC]^10^ or carcinoid tumors^11^), as well as tumors that bear some histologic resemblance to SCLC (e.g. SMARCA4-deficient undifferentiated tumor^12^), it does not in treatment-naïve SCLC tumor samples^4, 13^. Nevertheless, the detection of YAP1 expression in SCLC patient-derived models^4, 9^ and previous evidence in SCLC genetically-engineered mouse models (GEMMs) of the temporal emergence of non-neuroendocrine, YAP1-positive states^14, 15^ suggests that YAP1 may indeed play a critical, if not subtype-distinguishing, role in SCLC biology. However, obtaining longitudinal patient biopsies, in particular having one collected prior to any treatment and another following relapse, is incredibly rare, and greatly limits our knowledge of common resistance mechanisms in SCLC. This is evidenced by the paucity of published examples of paired transcriptional analyses^16, 17^. Additionally, surgical resections are not performed commonly in SCLC patients and core needle biopsies only provide one or two small core samples for research, resulting in an inability to conduct numerous analyses.

In this report, we find that YAP1 expression in treatment-naïve SCLC is rare and negligible, while single-cell analyses and IHC from relapsed SCLC patient tumors and methylation analyses of circulating tumor (ct)DNA and mass cytometry from circulating tumor cells (CTCs) from relapsed patients collectively indicate an emergent, YAP1-positive cell population that coincides with treatment resistance. These nascent YAP1(+) cells are neuroendocrine-low, mesenchymal, senescent, and transcriptionally plastic. Paired analyses indicate that while YAP1(+) cells emerge late in the real-time natural history of the SCLC course, they represent evolutionarily ancestral cells (i.e. stem-like pre-cursors) in pseudo-time, suggesting the YAP1(+) cells represent at stem-like source for relapsing tumor.

Surprisingly, YAP1(+) cells’ transcriptional profiles most closely resemble LCNEC rather than SCLC and, as a result, possess a unique surface proteome, absent traditional neuroendocrine-high SCLC cell surface targets such as Delta-like ligand 3 (DLL3) and Seizure-related homolog 6 (SEZ6), while exhibiting such candidate surface targets as Trophoblast cell surface antigen 2 (TROP2) and B7 Homolog 3 (B7-H3) previously highlighted in YAP1-positive LCNECs^10^.

Collectively, these findings demonstrate that YAP1 expression, while not a subtype-defining feature of treatment-naïve SCLC, is an inducible feature of relapsed SCLC, underlying its recalcitrant biology. This observation, in turn, reconciles the apparently conflicting accounts of YAP1 in SCLC and highlights a putative mechanism for several consistent observations in SCLC, including diminishing neuroendocrine character at relapse and frequent mixed LCNEC/SCLC histology. Most importantly, the identification of these YAP1(+) cells offers a critical population of intractable cells present only in the relapsed state against which to direct therapeutic development, including with cell surface targeting strategies.

## Results

### YAP1 induction following platinum chemotherapy

While YAP1 was undetectable in treatment-naïve SCLC and most SCLC cell lines, cisplatin-resistant cells (H69/CPR) had higher levels of YAP1 by reverse phase protein array (RPPA) compared to parental H69 cells and untreated cell line xenograft tumors (Fig. 1a). Similarly, SCLC-A cell lines treated with cisplatin for five days demonstrated increased YAP1 and NOTCH2 levels, along with decreased ASCL1. (Fig. 1b). Similarly, RNAseq analysis of biopsies from treatment naïve and relapsed SCLC patients in two independent datasets^16, 17^ revealed higher *YAP1* expression and YAP1 pathway activation, indicated by the YAP/TAZ target score^18^, at relapse (Fig. 1c; Supplemental Fig. 1a). This score consists of 19 genes and was established using over 9,000 specimens from over 30 different tumor types^18^.

**Fig. 1:**
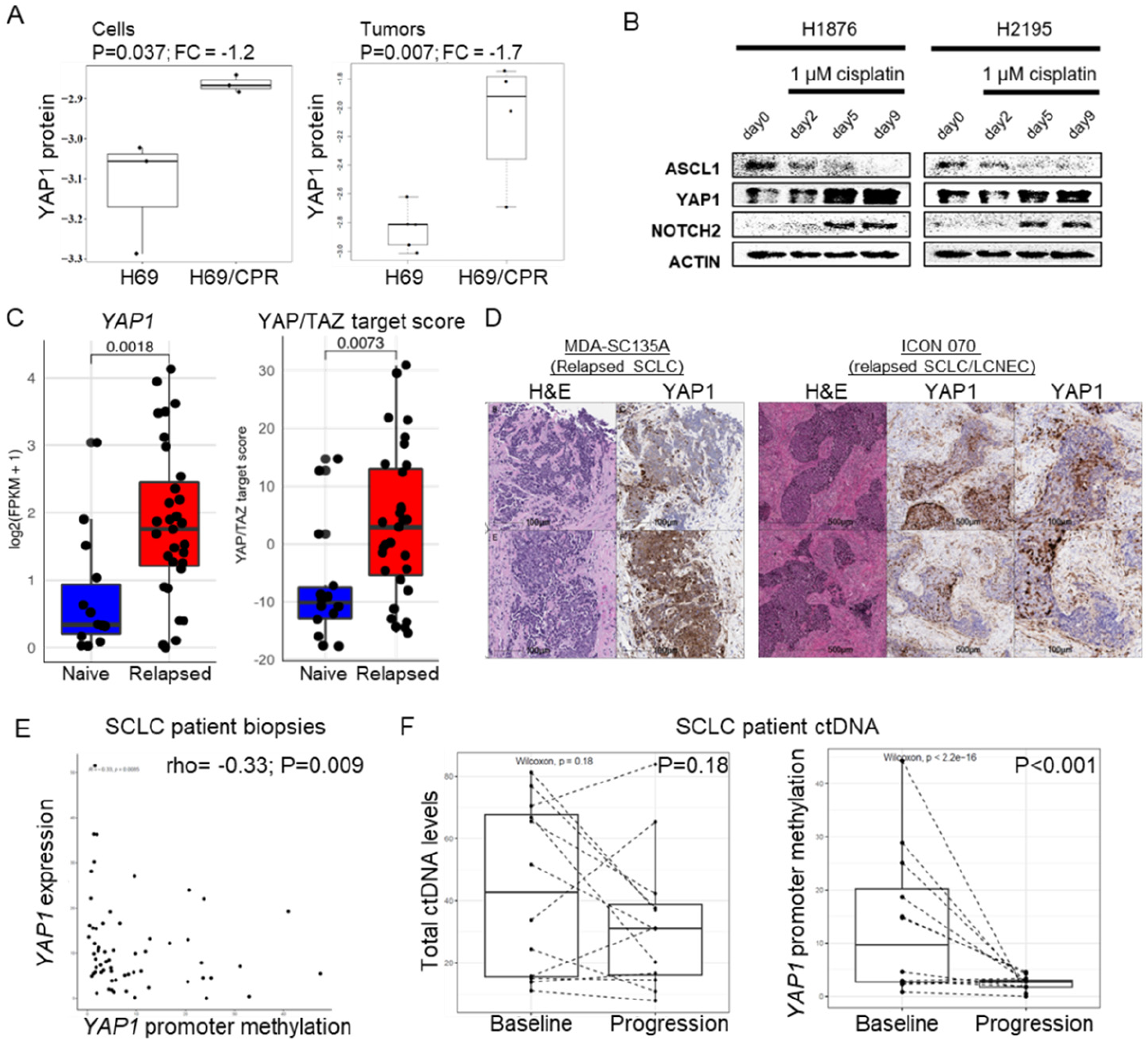
YAP1 is virtually undetectable in treatment naïve SCLC, but is upregulated following relapse to chemotherapy. A, YAP1 protein levels in H69 and H69/CPR cells and tumors following cisplatin treatment by RPPA. B, Extended treatment of SCLC cell lines with cisplatin decreases ASCL1 and increases YAP1 and NOTCH2 levels. C, Bulk RNAseq analysis of *YAP1* (left) and YAP/TAZ target score in SCLC patient tumors at treatment naïve and relapsed timepoints. D, Representative IHC demonstrating nuclear YAP1 in SCLC cells from a relapsed pure SCLC tumor (left) and in a mixed histology tumor (right). Scale bar = 100 or 500 μm. G, *YAP1* promoter methylation correlates with *YAP1* expression in SCLC patient biopsies. H, ctDNA levels are unchanged between baseline and relapsed samples (left), but *YAP1* promoter methylation is detectable only in baseline samples (right).

### YAP1 in relapsed SCLC

To determine whether nuclear YAP1 was present in histologically-confirmed SCLC cells from relapsed SCLC patients, YAP1 IHC was performed. In contrast to treatment naïve SCLC samples, nuclear and cytoplasmic YAP1 was expressed by cells histologically consistent with SCLC both in relapsed pure SCLC biopsies and in mixed histology biopsies (Fig. 1d). As an example of acquired YAP1 being associated with progression, pre- and post-biopsy scans demonstrate tumor progression from YAP1-expressing patient SC135, but not from YAP1-negative patient SC142 (Supplementary Fig. 1b).

However, bulk transcriptome data does not allow for the assignment of *YAP1* expression to tumor cells, therefore the presence of YAP1-expression on CTCs was confirmed by mass cytometry, or CyTOF. SCLC patients have high numbers of CTCs, which represent metastatic cell populations and reflect burden of disease^19-21^. CTCs were identified as being positive for CD56, a commonly used neuroendocrine marker for hgNEC diagnosis, and negative for CD45 to exclude immune cells (Supplemental Fig. 1c). Mass cytometry identified CTCs from seven SCLC patients using a single vial of blood from each. Additionally, phosphoYAP1 (Y357) levels were detectable in CTCs from all patients (Supplementary Fig. 1d). Specifically, one patient exhibited increased phosphoYAP1 and vimentin, a mesenchymal protein, levels in the second of two longitudinal samples (Supplemental Fig. 1d-e). This data demonstrates a trend for higher phosphoYAP1 levels in most relapsed patients, compared to the treatment naïve sample. PhosphoYAP1 levels detected in patient liquid biopsies were consistent with single-cell RNAseq detection of *YAP1* expression in tissue biopsies from the same patients (Supplemental Fig. 1f,g).

Epigenetic regulation of YAP1 via promoter methylation has been described previously in another pulmonary high-grade neuroendocrine carcinoma^10^, as well as breast cancer^22^. *YAP1* expression was negatively correlated with promoter methylation in SCLC patient biopsies (Fig. 1e) and cell lines (Supplemental Fig. 1h). While total ctDNA levels were unchanged between baseline and progression samples from SCLC patients (Fig. 1f, left), *YAP1* promoter methylation was not detectable at progression. Paired ctDNA samples also demonstrated abundant *YAP1* promoter methylation at baseline, followed by repression of methylation at progression (Fig. 1f, right).

### Single-cell transcriptional profiling of relapsed SCLC

To investigate inter- and intra-tumoral heterogeneity, core needle biopsies, endobronchial ultrasound (EBUS) biopsies, and surgical resections from previously treated SCLC (n=15) patients were dissociated and analyzed by single-cell RNAseq (Supplemental Table 1-2; Supplemental Fig. 2). A total of 69,677 cells passed the initial quality control analysis and were classified by cell type (Fig. 2a). The 35,920 pooled cancer cells were selected based on cell calls, expression of neuroendocrine genes, and inferred copy number analysis (Supplemental Fig. 3a). Cancer cells were visualized before debatching to determine intertumoral heterogeneity (Supplemental Fig. 3b), but all analyses were performed on debatched samples. When cancer cells were color-coded by biopsy/resection site (i.e., liver, lymph node, adrenal, lung, etc.), then no clear transcriptional patterns were present (data not shown, biopsy site listed in Supplemental Table 1). Similarly, cancer cells were color-coded by patient without strongly defined clusters by individual, indicating inter-tumoral heterogeneity (Fig. 2b, upper left), and suggesting that common transcriptional pathways/mechanisms are present in biopsies from distinct patients. In fact, when the eight cell clusters from a single patient biopsy (MDA-SC135A) were highlighted, they were located throughout the pooled cancer cell populations (Supplemental Fig. 3c).

**Fig. 2:**
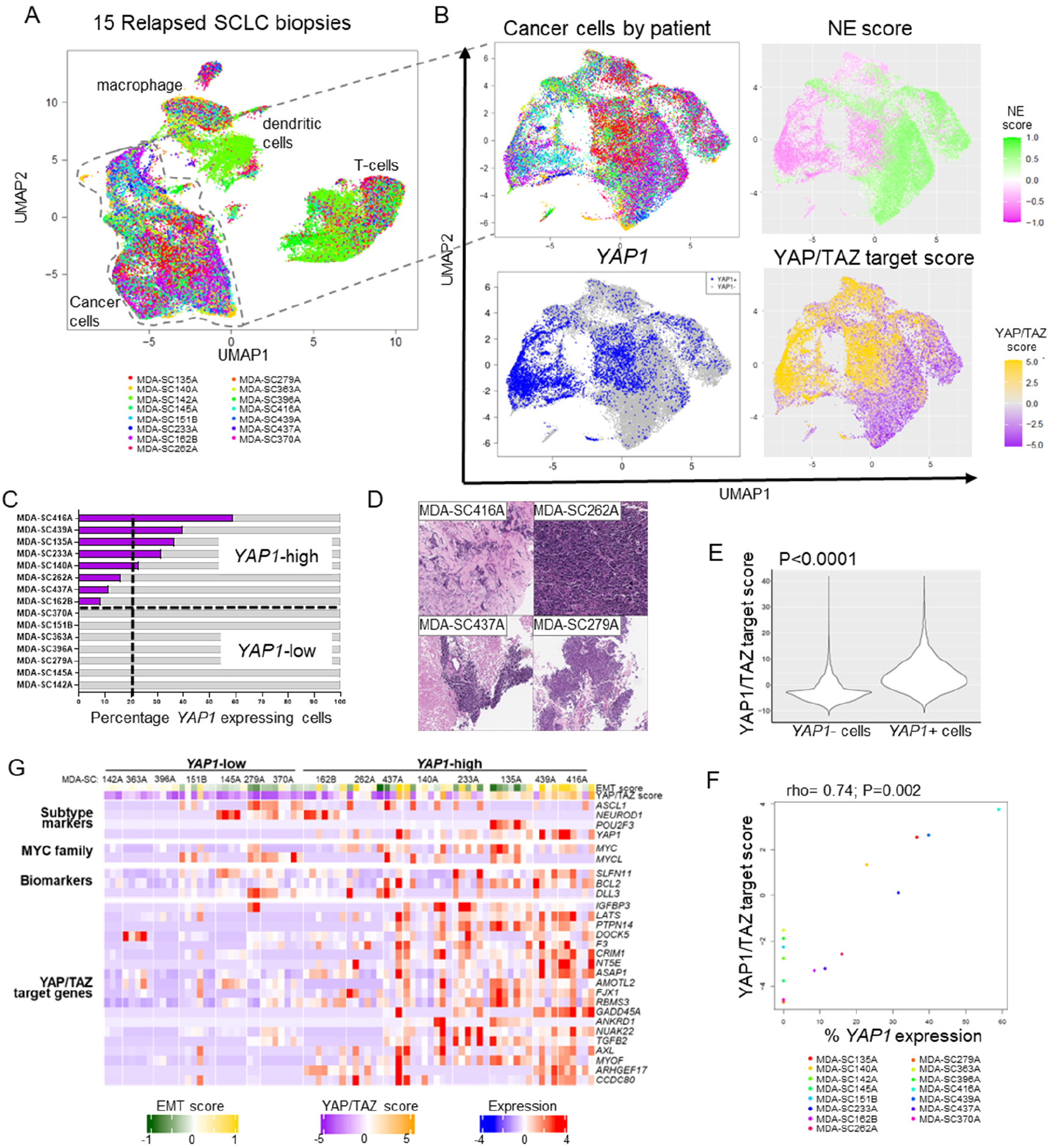
YAP1 is detectable in relapsed SCLC, even when absent at diagnosis. A, UMAP plot of all cell populations from 15 SCLC biopsies collected from relapsed patients. B, UMAP plots of cancer cell populations reveal *YAP1-*expressing cell populations with elevated YAP/TAZ target score and low NE score. C, Percentage of *YAP1*(+) cells in each biopsy. Broken line indicates the cut-off between *YAP1*-high and -low status. D, SCLC morphology of additional cores from YAP1-high biopsies. E, *YAP1*(+) cells have a higher YAP/TAZ target score. F, Percent YAP1 expression correlates with YAP/TAZ target score in patient biopsies. G, Heatmap demonstrating individual cluster expression from each relapsed biopsy sorted by YAP1 status including expression of EMT and NE scores, subtype genes, MYC family members, SCLC biomarkers, and YAP/TAZ target genes.

To determine expression of common markers of SCLC in the pooled cancer cells, binary gene expression or molecular scores were visualized on UMAP plots. YAP1 was virtually undetectable in bulk analyses of treatment naïve SCLC tumors^10^. However, in relapsed SCLC biopsies, *YAP1* was expressed by 15.34% of cancer cells in the pooled population from 15 patients and 0 to 59.06% cells in individual relapsed patient biopsies (Fig. 2b, bottom left and Supplemental Table 3). As expected, *YAP1* expression occurs in cell populations with a low NE score (Fig. 2b, upper right) and high YAP/TAZ target score (Fig. 2b, bottom right). For comparisons of YAP1-high vs. YAP1-low biopsies, biopsies with any detectable YAP1 expression was used as the cut-off for defining these groups (Fig. 2c). Additional cores or surgical resections collected at the same time point were analyzed histologically by thoracic pathologists for morphological changes. All specimens analyzed were determined to be morphologically consistent with SCLC (Fig. 2d). As expected, *YAP1* mRNA expression was higher in the YAP1-high biopsies and *YAP1(+)* cells (P<0.0001 for both; Supplemental Fig. 3d) and the YAP/TAZ target score was higher in *YAP1(+)* cells (P<0.0001; Fig. 2e). In fact, mean YAP/TAZ target score correlated directly with YAP1 percentage (Fig. 2f).

### Transcriptional diversity in relapsed SCLC

In order to determine specific differences in YAP1(+) and (-) cells, gene set enrichment analysis (GSEA) was performed. There was enrichment of TNF alpha, EMT, TGF beta, and multiple inflammatory pathways in *YAP1*(+) cells (Supplemental Fig. 2e). While some individual genes appeared to be abundant in YAP1-high biopsies and in *YAP1*(+) cells (e.g., *YAP1, MALAT1, FTH1, KRT17, CXCL8, ANXA1, S100A6, FTL, MT2A*, etc.), significant transcriptional variation was observed across cellular subpopulations, or clusters. Therefore, gene expression of molecular subtypes (*YAP1, ASCL1, NEUROD1, POU2F3*), mesenchymal EMT genes (*ZEB1, ZEB2, SNAI2, VIM*), epithelial genes (*EPCAM, CDH1*), NE genes (*INSM1, SYP, UCHL1*), non-NE genes (*REST, ANXA1*), biomarkers (*SLFN11, BCL2, DLL3*) and YAP/TAZ target score genes were determined in YAP1-high and -low patient biopsies (Fig. 2g). Similar to SCLC patient derived xenograft models^20^, significant variation in gene expression occurred across clusters in relapsed SCLC patient biopsies, regardless of YAP1 status. A greater number of clusters express NE genes in YAP1-low biopsies. There were no differences in cluster numbers present in individual biopsies between YAP1-low and -high biopsies (Supplemental Table 1; p=0.25). In SCLC PDX models, treatment naïve tumors had fewer clusters than relapsed tumors^20^, suggesting that this may be an indicator of intratumoral heterogeneity (ITH). Similarly, GSEA analysis of YAP1-high and -low biopsies revealed higher inflammatory response pathways, TNF alpha and TGFB signaling, and EMT in the YAP1-high relapsed biopsies (Supplemental Fig 3f).

Genomic loss of *CDKN2A* or *SMARCA4* frequently occur in YAP1-high LCNEC^10^ and YAP1 has been correlated with SMARCA4-deficient undifferentiated tumors in specimens initially identified as SCLC^12^. To further ensure that *YAP1(+)* cells or clusters with *YAP1(+)* cells were SCLC, *CDKN2A* and *SMARCA4* were evaluated. The left cluster with the most *YAP1(+)* cells had low expression of both *CDKN2A* and *SMARCA4* (Supplemental Fig. 3g). Along with thoracic pathology analysis of biopsies, this data suggests that these cells were indeed SCLC and do not represent SMARCA4-deficient carcinomas.

### SCLC molecular subtype markers in relapsed disease

Our group^4^ and others have previously defined SCLC subtypes and expression of the transcription factors defining these groups was investigated in the relapsed patient biopsies. As expected, *ASCL1, NEUROD1*, and *POU2F3* were expressed by relapsed SCLC cells (Fig. 3a; Supplemental Table 3). *NEUROD1* and *YAP1* were expressed in a single biopsy (MDA-SC162A), but there was overlap between *ASCL1* or *POU2F3* with *YAP1* (Fig. 3b). This is consistent with previous reports^23^. While all molecular subtypes were observed in the 15 relapsed biopsies, primarily SCLC-A and SCLC-P biopsies had *YAP1(+)* cells (Fig. 3b and Supplemental Table 4). Notably, none of the SCLC-I biopsies contain *YAP1*(+) cell populations. Consistent with SCLC-A biopsies having *YAP1*(+) populations, *YAP1* expression was higher in relapsed SCLC-A subtype patients (Fig. 3c). While the numbers are small, a similar pattern is observed in SCLC-N patients (Supplemental Fig. 4a). In the absence of having performed single-cell RNAseq on diagnosis biopsies, retrospective analysis was performed by IHC to determine whether YAP1 expression was measurable prior to any therapy. Biopsies that were YAP1-high at relapse (by single-cell RNAseq (Fig. 2c) had no detectable YAP1 by IHC at the time of diagnosis (i.e., prior to treatment), but similar ASCL1 levels (Fig. 3d).

**Fig. 3:**
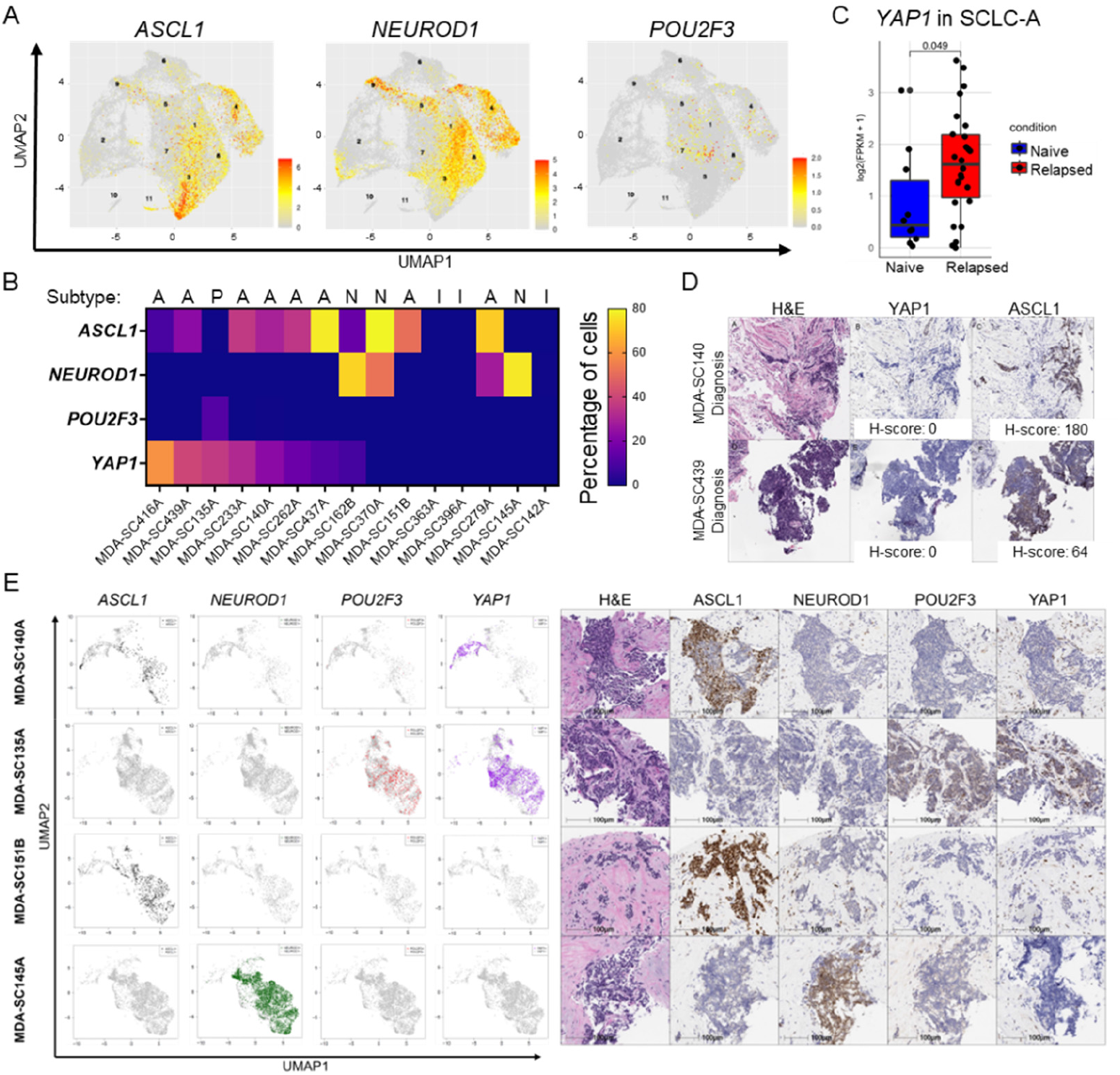
YAP1 in relapsed SCLC is mutually exclusive with ASCL1 and NEUROD1. A, UMAP feature plots demonstrating expression levels of *ASCL1, NEUROD1*, and *POU2F3* in pooled, relapsed SCLC biopsies. Cluster numbers are indicated on the plots. B, Heatmap sorted by *YAP1* expression demonstrating molecular subtype and co-expression of transcription factors. SCLC molecular subtype is listed at the top. C, Bulk RNAseq analysis of *YAP1* in SCLC-A subtype patient tumors at treatment naïve and relapsed timepoints. D, Paired biopsies from patients that are *YAP1*-high at relapse have undetectable YAP1 at diagnosis by IHC. E, Single-cell RNAseq and IHC analysis of core needle biopsies collected at the same time. Scale = 100 μm.

Five of the biopsies had both IHC and single-cell RNAseq for ASCL1, NEUROD1, POU2F3, and YAP1 from separate cores collected at the same time point. Data was concordant between both techniques for all but one biopsy (MDA-SC140A; Fig. 3d, Supplemental Table 3). This biopsy had 22.81% *YAP1(+)* cells by single-cell RNAseq and no detectable YAP1 by IHC, suggesting that 1) there may be variation between the paired core biopsies analyzed; 2) the single-cell RNAseq is more accurate at detecting expression in fewer than 25% of cells; or 3) that YAP1 is regulated post-transcriptionally. To exclude the possibility that the single-cell RNAseq data was detecting *YAP1* expression in stromal cells, expression of neuroendocrine genes (e.g., *UCHL1*) was verified in *YAP1*(+) cell populations, albeit at lower levels that are not depicted in the binary feature plots (Supplemental Fig. 4b). In order to determine if gene expression was mutually exclusive, expression of *YAP1* in *ASCL1, NEUROD1*, or *POU2F3* positive and negative cells was determined. *YAP1* was not expressed by *ASCL1* or *NEUROD1* positive cells, confirming that these genes were not co-expressed by the same cells (Supplemental Fig. 4c). As expected, *NEUROD1* was expressed equally in *ASCL1* positive and negative cells (Supplemental Fig. 4c). Along the same lines, *YAP1* was expressed by *POU2F3* positive cells by transcriptional profiling and also by IHC in biopsy MDA-SC135A. Interestingly, EMT score demonstrated a wide range, in spite of *YAP1* status, suggesting that there were both *YAP1*(+) epithelial and *YAP1*(+) mesenchymal cells present in relapsed patient biopsies (Supplemental Fig 4d). This is likely due to *YAP1*(+) cells undergoing partial EMT or the population co-expressing *POU2F3* having a lower EMT score and being more epithelial (Supplemental Fig. 4e).

### Transcriptional Plasticity

Tumors were ranked by YAP1% cells and EMT score^24^, NE score^25^, ITH score^20^, and CytoTRACE analysis to predict differentiation state of cells was performed (Supplemental Fig. 5a; cytotrace.stanford.edu). High CytoTRACE scores, associated with low differentiation, can additionally predict the presence of cancer stem cell populations^26^. There was a broad range of values for EMT score, NE score and CytoTRACE values within individual samples, suggesting significant heterogeneity within each sample.

Cell transport potential, a measure of plasticity^27, 28^, was calculated for the cancer cells in 15 relapsed patient biopsies. In general, cell transport potential was high in nearly all cell populations with a smaller range than in previous datasets (Fig. 4a)^4, 27^. The velocity streams indicate the RNA velocity originates from two distinct regions (left and right). The velocity streams originate from the *YAP1(+)* regions when they are overlayed with binary expression of *YAP1* (Fig. 4b). This is additionally confirmed with Partition-Based Graphical Abstraction (PAGA), a computational tool that classifies clusters chronologically based on their continuous cell transitions inferred from RNA velocity (Fig. 4c). Additionally, average cell transport potential was higher in *YAP1*(+) compared to *YAP1*(-) cell populations (Fig. 4d), resulting in a shift of peaks (Supplemental Fig. 5b).

**Fig. 4:**
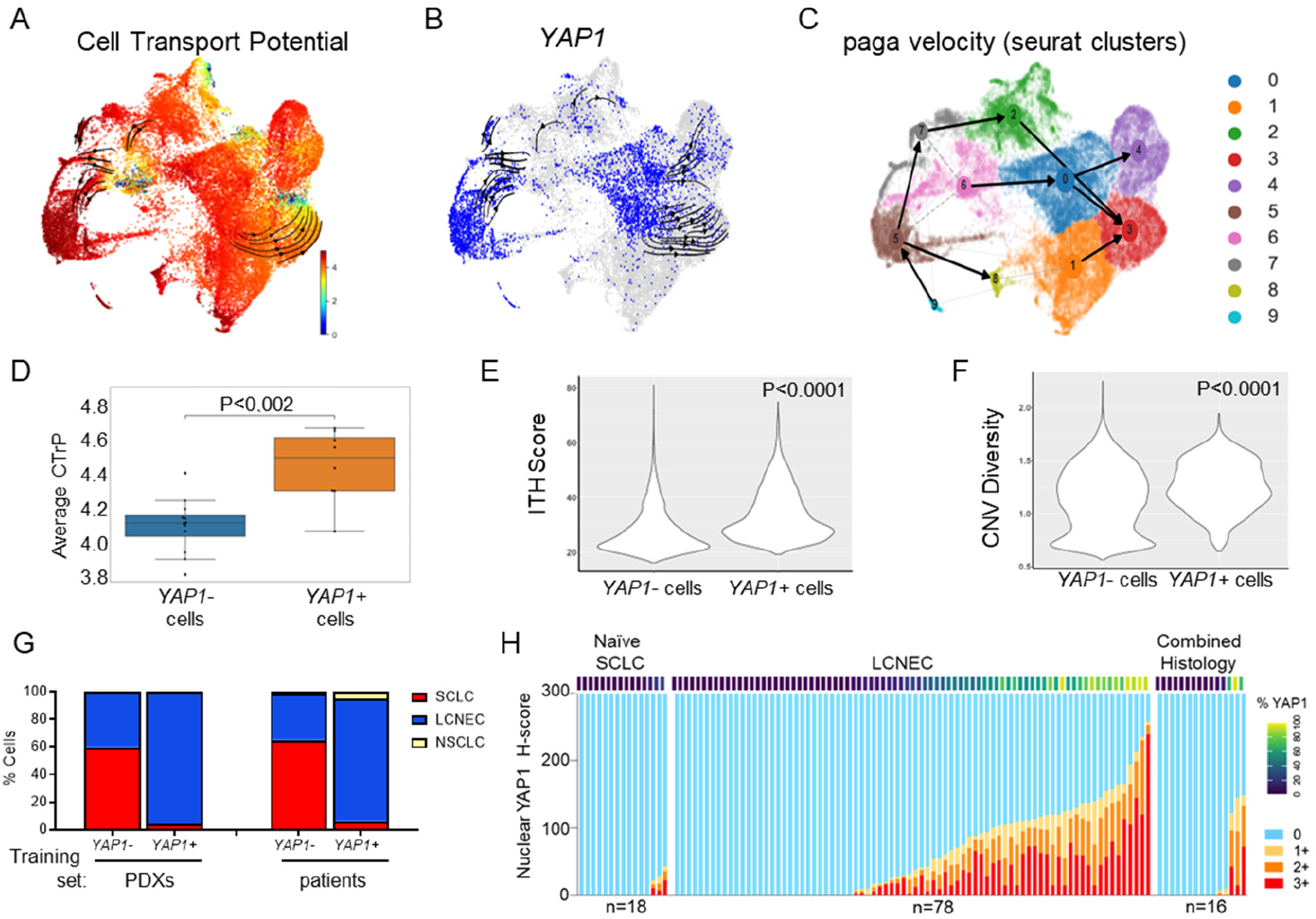
YAP1 in relapsed SCLC is associated with EMT, senescence, and plasticity. A, Cell transport potential (CTrP) in relapsed biopsies. B, RNA velocity vector streams originating from *YAP1*(+) populations. C, PAGA velocity maps demonstrate origins of arrows from *YAP1*(+) clusters. D, Average CTrP of *YAP1*(+) and (-) populations demonstrating higher overall plasticity in *YAP1*(+) cells. E, Comparison of ITH score in *YAP1*(+) and (-) populations. F, Analysis of CNV diversity in *YAP1*(+) and (-) cells. G, Cell type annotation of *YAP1*(+) and *YAP1*(-) cells using two independent training sets, PDX (left) and patient biopsies (right). H, Nuclear YAP1 H-score values and percentage in SCLC, LCNEC, and combined histology tumors.

Consistent with this, cells from *YAP1*-high biopsies and *YAP1*(+) cells demonstrate an increased ITH score compared to those in the *YAP1*-low biopsies and *YAP1*(-) cells and there was a direct correlation between percent YAP1(+) cells and ITH score (Fig. 4e, Supplemental Fig 5c,d). Similarly, chromosomal instability is a hallmark of cancer that is elevated in SCLC^29^ and associated with reduced prognosis, therapeutic resistance, cell cycle arrest, and senescence^29-31^. Chomosomal instability was measured using diversity of copy number variation inferred from single-cell RNAseq data^30^ and *YAP1*(+) cells exhibited elevated copy number variation (CNV) diversity, suggesting increased chromosomal instability (Fig. 4f).

### Histology of YAP1(+) cell populations

Evaluation of YAP1 IHC by thoracic pathologists (Fig. 1d, 3e) revealed nuclear YAP1 in SCLC cells from relapsed patients; however SCLC and LCNEC are notoriously difficult to distinguish histologically. In order to determine transcriptional histology of cell populations in relapsed SCLC, either PDXs or patient biopsies with verified histology confirmation were used as training sets in Cell Typist (celltypist.org), an open source platform to define cell type annotations using single-cell RNAseq data^32, 33^, to transcriptionally distinguish SCLC, LCNEC, and NSCLC cells. Both training sets determined that approximately 60% of *YAP1*(-) cells were SCLC, while less than 5% of *YAP1*(+) cells were SCLC (Fig. 4g). The majority of *YAP1*(+) cells were determined to be LCNEC, rather than NSCLC. Consistent with this, roughly half of LCNEC tumors had detectable nuclear YAP1^10^ (Fig. 4h). This suggests that the YAP1(+) cells in relapsed SCLC biopsies may indicate a transcriptional LCNEC-like reprogramming and histologic plasticity.

### Pseudotime analysis of relapsed SCLC

In order to investigate lineage of relapsed cell populations, pseudotime trajectory analysis using CellRank2^34^ was performed. Patient MDA-SC162 received two biopsies, the first (A) following frontline chemo-immunotherapy and the second ten months later following progression on maintenance immunotherapy (B) (Fig. 5a). Clustering of cells reveals many common clusters containing cells from both biopsies (Fig. 5b). Pseudotime trajectory analysis of these biopsies demonstrated three monocle states, two of which contained cells from both biopsies (upper and left) and one state (right) with cells primarily from the second biopsy (Fig. 5c; left). Notably, the earliest pseudotime occurs in the state consisting of cells from the second biopsy, suggesting that highly plastic cells are present following disease progression (Fig. 5c; right). Accordingly, *ASCL1*- and *NEUROD1*-positive cells are located along this same monocle state, with *YAP1* at the far tip.

**Fig. 5:**
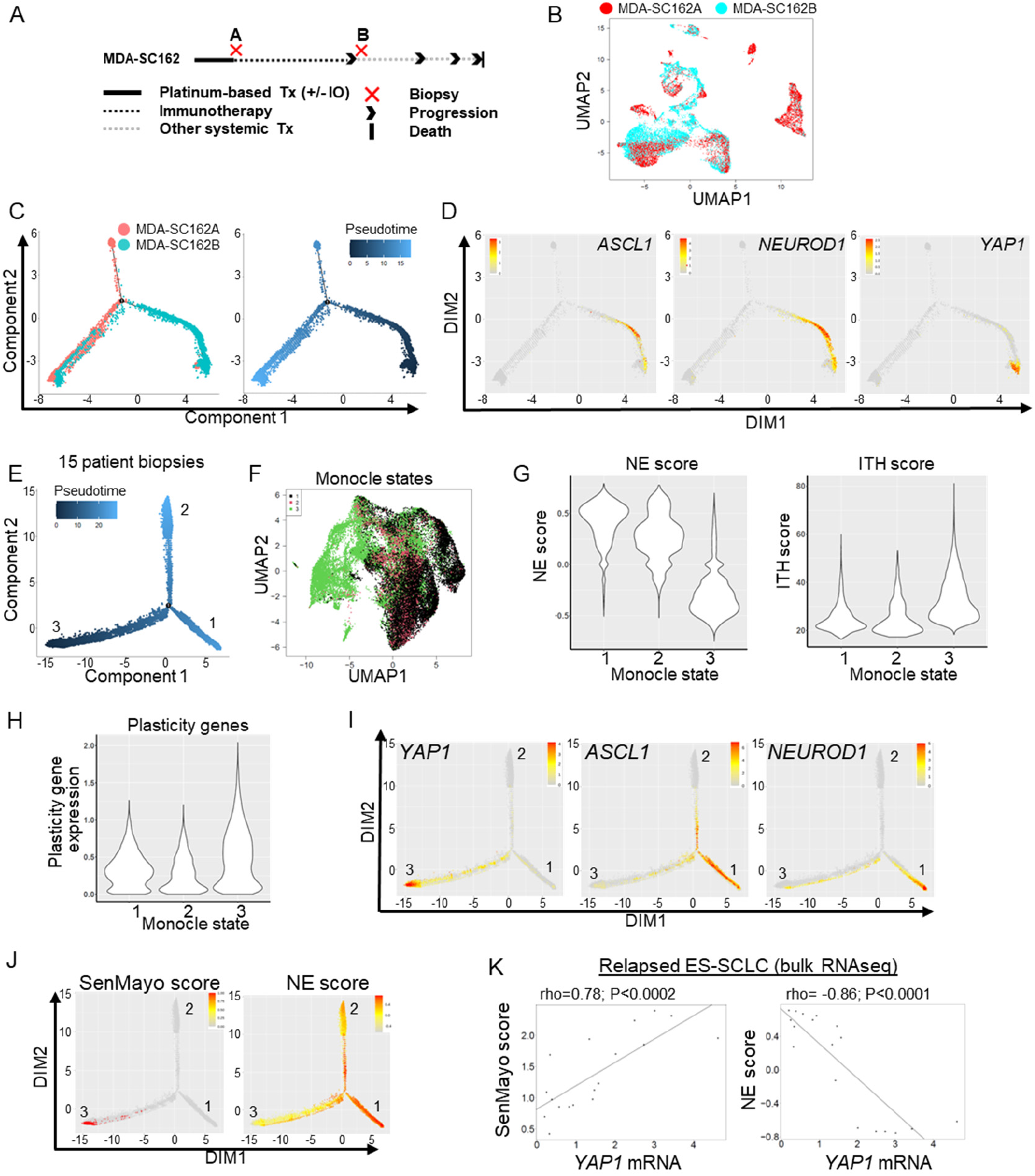
Pseudotime trajectory of relapsed SCLC biopsies. A, Patient MDA-SC162 treatment course, including timing of paired biopsies. B, UMAP plot of biopsies MDA-SC162A and MDA-SC162B. C, Pseudotime trajectory analysis of paired MDA-SC162 biopsies colored by biopsy (left) and pseudotime (right). D, Analysis of paired MDA-SC162 biopsy cells *ASCL1, NEUROD1*, and *YAP1* expression in pseudotime. E, Pseudotime trajectory analysis of 15 pooled patient biopsy cancer cells. F, Monocle states plotted in the UMAP space. G,H, Abundance of NE score, ITH score (G), and plasticity genes (H) in the monocle states. I, Analysis of *YAP1, ASCL1*, and *NEUROD1* expression in pseudotime. J, SenMayo and NE scores in pseudotime. K, Correlation between SenMayo or NE score and *YAP1* expression in bulk relapsed SCLC tumors.

Analysis of all samples pooled together revealed three, roughly equal monocle states, or lineages. Similar to MDA-SC162 biopsies, a single monocle state (state 3) exhibits the earliest pseudotime (Fig. 5e). When monocle states are projected into UMAP space, state 3 is distinct from states 1 and 2 (Fig. 5f) and represent the cells primarily associated with low-NE scores and elevated ITH (Fig. 5g). While no difference was detected in entropy between monocle states (Supplemental Fig. 6a), there was an abundance of plasticity genes specifically in state 3, both for the cumulative score (Fig. 5h) and in the individual genes (Supplemental Fig. 6b). Consistent with the transcriptional plasticity of monocle state 3, both EMT and ITH scores are elevated in this particular branch (Supplemental Fig. 6c). Further, state 1 was associated with *ASCL1* and *NEUROD1* and state 3 was associated with *YAP1* expression (Fig. 5i; Supplemental Fig. 6c); which is consistent in both the dimensional (DIM) and UMAP space, suggesting that these two states are distant in pseuodtime. This distance between *YAP1* and *ASCL1* states is maintained across individual samples (Supplemental Fig. 6f).

To determine which cells were associated with senescence-associated pathways or senescence, the SenMayo score^35^ was analyzed in pseudotime. Similar to *YAP1*, SenMayo score was localized in state 3 (Fig. 5j, left) and *YAP1* percentage directly correlated with SenMayo score (Supplemental Fig. 6g). In contrast, NE score^25^ was localized primarily in states 1 and 2 (Fig. 5j, right) and was negatively correlated with *YAP1* percentage (Supplemental Fig. 6g). Despite relatively small numbers of cells expressing *YAP1* in relapsed disease, bulk analyses of patient tumors^36^ also demonstrated that *YAP1* was correlated with SenMayo score^35^, while negatively correlated with NE score^25^ (Fig. 5k). Collectively, this data suggests that the *YAP1*(+) cell populations have undergone senescence and are transcriptionally diverse.

### YAP1 and MYC in Relapsed SCLC

Previous reports have demonstrated that MYC is a resistance mechanism in SCLC,^37, 38^ driving dedifferentiation of tumor cell state towards YAP1^15^. Therefore, we investigated the link between YAP1 and MYC in relapsed SCLC patient biopsies. *YAP1* and *MYC* were expressed by similar populations (clusters 1 and 2; Fig. 6a; Supplemental Table 3). However, only cluster 1 (left side of the UMAP plot) had high EMT score (Fig. 6a, right panel). In the 15 relapsed biopsies, average cell transport potential was not different between *MYC(+)* and *MYC(-)* populations (Supplemental Fig. 7a), despite frequent co-expression of both genes (Fig. 6b) and higher *YAP1* expression in *MYC(+)* cells (Fig. 6c). Similarly, YAP1 and MYC were co-expressed by IHC in PDX tumors derived from patients with relapsed disease, in the absence of genomic or extrachromosomal amplifications (Fig. 6d). Similar to *YAP1, MYC* is expressed in monocle state 3. However, *MYC* is localized both at the tip and closer to the central branch point (Fig. 6e). Notably, this state localization is common to *REST, SOX9*, and *SNAI1* (Fig. 6e).

**Fig. 6:**
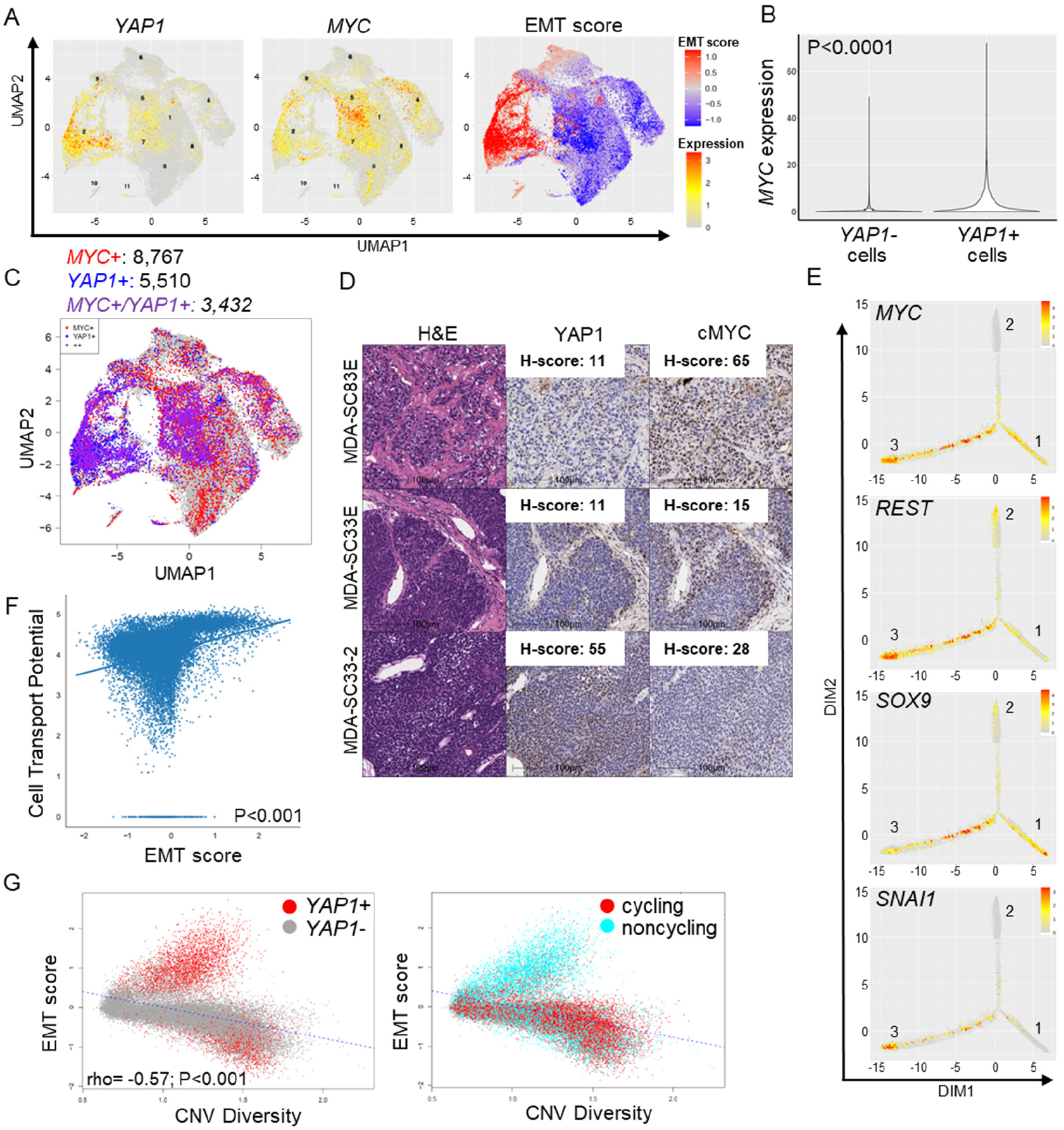
EMT is associated with plasticity and *YAP1* and *MYC* are co-expressed. A, UMAP feature plots demonstrating *YAP1, MYC*, and EMT score in relapsed SCLC biopsies. Cluster numbers are indicated on the plots. B, Violin plots demonstrating *MYC* expression in *YAP1*(+) cells. C, *YAP1* and *MYC* are frequently co-expressed in relapsed SCLC biopsies. D, IHC for YAP1 and MYC with positive expression of both in PDX tumors derived from relapsed SCLC patients. Scale = 100 μm. E, Visualization of *MYC, REST, SOX9*, and *SNAI1* in pseudotime demonstrating similar expression patterns. F, CTrP is directly correlated with EMT score. G, EMT score correlation with CNV diversity demonstrates a different pattern for epithelial and mesenchymal cells (left) and that *YAP1*(+), noncycling cells correlate with chromosomal instability (right).

### YAP1 and EMT are associated with chromosomal instability

EMT is a known mechanism of cancer resistance. Therefore, the hypothesis that EMT is associated with plasticity and chromosomal instability in relapsed SCLC biopsies was tested. *YAP1* expression also correlates with EMT in both relapsed SCLC (Supplemental Fig. 7b) and LCNEC^10^. Similar to cell transport potential being elevated in *YAP1*(+) cells, cell transport potential was positively correlated with EMT score (Fig. 6f). Chromosomal instability can indirectly induce EMT^39^. Consistent with this, when EMT score was correlated with chromosomal instability, the *YAP1*(+) mesenchymal cells were positively correlated and epithelial cells were negatively correlated with inferred CNV diversity (Fig. 6g, left). Furthermore, these *YAP1*(+) mesenchymal cells were largely non-cycling (Fig. 6g, right), which is consistent with the correlation between *YAP1* and the SenMayo score (Fig. 5j,k; Supplemental Fig. 6e).

### Cell Surface Targets in relapsed SCLC

Several emerging anti-cancer strategies exploit the tumor cell surfaceome to deliver cytotoxics (antibody-drug conjugates [ADCs]) or immune cells (T-cell engagers [TCEs] or chimeric antigen receptors [CARs]). While tarlatamab (DLL3-TCE) is currently the only FDA-approved surface targeting strategy in relapsed SCLC^40, 41^, several have been explored and shown activity--including ADCs (SEZ6 [ABBV-706^42^]; B7-H3 [ifinatamab deruxtecan^43^, HS-20093/GSK5764227^44^]; and TROP2 [sacituzumab govitecan^45^, SHR-A1921^46^) bispecific TCEs (DLL3; BI764532^47^), trispecific TCEs (HPN328^48^, RO7616789/RG6524^49^), and DLL3 CAR-Ts (AMG119^50^, LB2102^51^). Consistent with this, surface target gene expression is abundant in bulk analysis of relapsed SCLC biopsies (Fig. 7a). Specifically, DLL3 is expressed in both treatment naïve and relapsed SCLC from the same patient (Fig. 7b); although expression is heterogeneous at relapse^20^. Additionally, expression of surface targets may occur in distinct cell populations. While DLL3 and SEZ6 are both expressed by similar cell populations, B7-H3 is expressed in distinct cell populations (Fig. 7c). Potential surface targets investigated in relation to *YAP1* expression include *CD276* (B7-H3), *TACSTD2* (TROP2), *ERBB2* (HER2), *DLL3, SEZ6*, and *CEACAM5*. All of these are being investigated for efficacy in preclinical studies or clinical trials. *CD276, TACSTD2*, and *ERBB2* are coexpressed by *YAP1*(+) cells (Fig. 7d; Supplemental Fig. 7c), with *CD276* expression occurring in both the left and right cluster of *YAP1* expressing cells. In contrast, *DLL3, SEZ6*, and *CEACAM5* are largely expressed by cell populations that are *YAP1(-)*. Pseudotime analysis of surface target expression reveals that *YAP1*-associated surface targets (*CD276, TACSTD2*, and *ERBB2*) are localized to monocle state 3, while the others (DLL3, SEZ6, and CEACAM5) are localized to monocle state 1 (Fig. 7e).

**Fig. 7:**
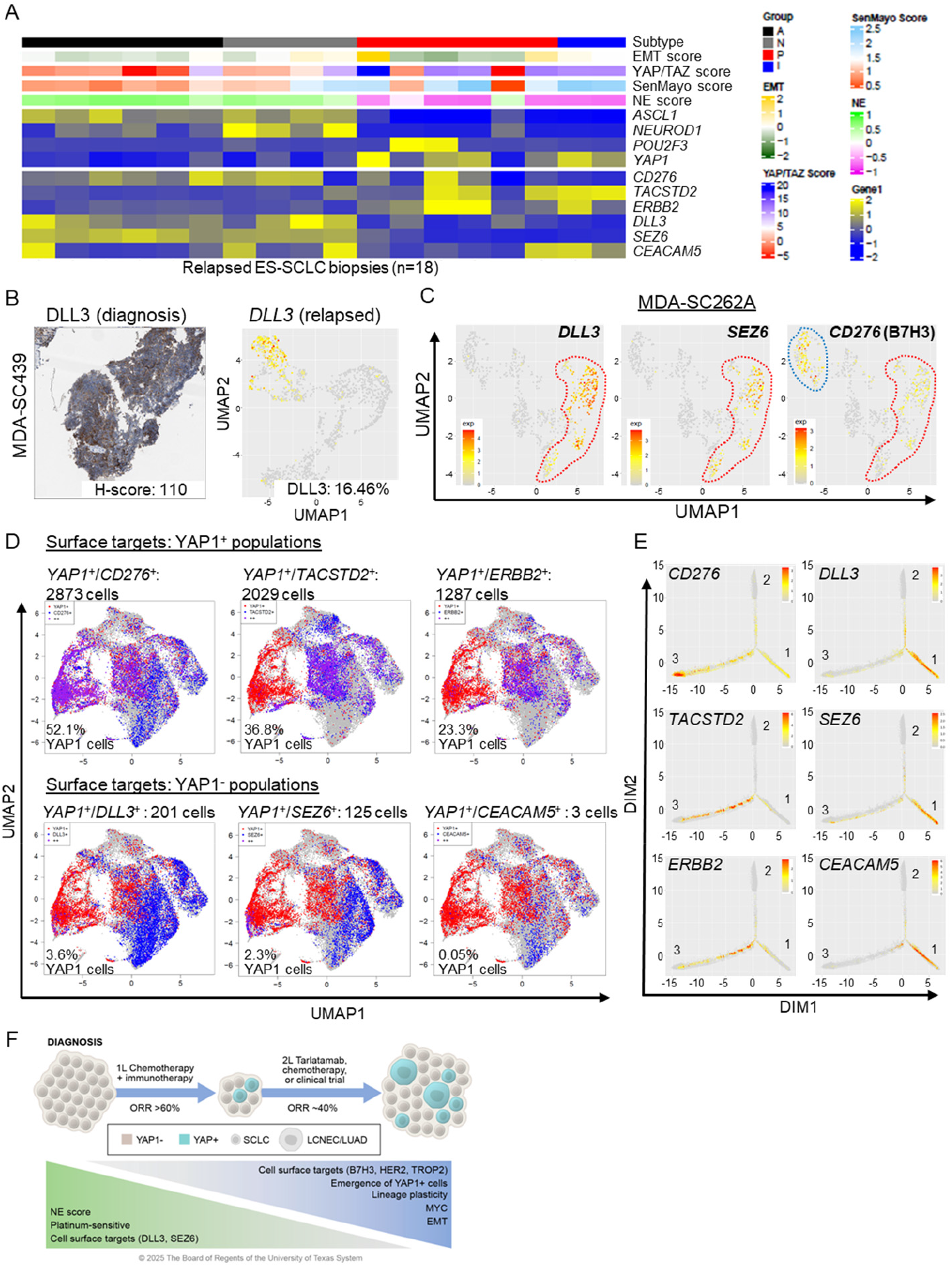
Cell surface proteome heterogeneity in relapsed disease may be harnessed with combination targeting. A, Heatmap demonstrating expression of molecular signatures, subtypes, subtype genes, and surface target genes in relapsed patient biopsies. B, DLL3 by IHC at diagnosis and by single-cell RNAseq at relapse in the same patient. C, UMAP feature plots demonstrating expression of surface targets in the same biopsy. *SEZ6* and *DLL3* are expressed in similar cell populations (red outline), but *CD276* (B7-H3) is expressed by a distinct population of cells (blue outline). D, UMAP plots with bicolor expression of *YAP1* (red) or cell surface targets (blue) to indicate co-expression of *CD276, TACSTD2*, and *ERBB2*, but not *DLL3, SEZ6*, or *CEACAM5*. E, DIM plots with expression of surface target genes in pseudotime space. F, Schematic demonstrating common features of diagnosis and relapsed SCLC biopsies.

In order to determine differences in immune cell populations, cell population make up was compared between *YAP1*-high and -low patients. Consistent with the increased inflammatory pathways associated with YAP1-high cancer cell populations, these same biopsies were enriched for cancer and NK cells (Supplemental Fig. 7d). Overall, YAP1 was not detectable in treatment-naïve pure SCLC, but YAP1(+) cells and increased transcriptional plasticity emerge following frontline platinum chemotherapy (Fig. 7f). Relapsed disease was associated with a reduction in NE score, platinum sensitivity and expression of cell surface targets, including DLL3 and SEZ6. In contrast, there was an expansion of YAP1 expression, EMT, plasticity and platinum resistance. However, cell surface targeting of these emerging populations exists via B7-H3 and TROP2 on the surface of *YAP1*(+) cells.

## Discussion

Both touted as a candidate subtype-defining feature and dismissed as a potential artifactual observation from stromal and/or mixed NSCLC contamination, recent years have seen considerable debate regarding the role of YAP1, if any, in SCLC. While frequent YAP1 expression in other neuroendocrine neoplasms (e.g. LCNEC, mixed histology neuroendocrine neoplasms, and carcinoid tumors) undoubtedly accounts for some of the confusion, the data presented here confirm that YAP1 is not only present, but biologically and clinically relevant, in relapsed SCLC.

Though nearly absent in the treatment-naïve setting of pure SCLC, YAP1 is consistently observed in both preclinical models and patient samples following treatment resistance and relapse. While treatment-naïve tumors are overwhelmingly defined by a single transcription factor and transcriptional subtype (e.g. ASCL1/SCLC-A), relapsed tumors instead emerge heterogeneously following treatment with YAP1 activation in a subset of cells at the expense of the previous dominant subtype-defining features. These tumors technically retain their original subtype identity in the bulk sense (i.e. they do not undergo complete subtype-switching), but subsequent treatments are now forced to grapple with the original tumor plus the nascent YAP1 population – a plastic, mesenchymal, non-neuroendocrine subset of cells markedly distinct from the remaining cells that resemble the treatment-naïve tumor. Indeed, pseudo-time analyses suggest that ASCL1 and YAP1 are present on distinct branches or lineages, despite being expressed within the same core biopsy.

Emergent YAP1(+) cells typically account for a minority of cells, even in the heavily pre-treated setting. However, the recalcitrant and highly plastic features of YAP1(+) cells are consistent with a potentially relentless subset of cells with both the ability to evade elimination by chemotherapy and immunotherapy, as well as the ability to replenish diminished populations of YAP1(-) cells to maintain tumor growth. This is especially evident in the pseudo-time analyses, including paired analyses, wherein *chronologically* later expression of YAP1 is shown to represent *evolutionarily* earlier events. In other words, the emergence of YAP1 is a de-differentation event toward a stem-like precursor that conceivably persist in the more quiescent *YAP1*(+) state to resist therapy pressure or replenish the more mitotically active neuroendocrine cells to support tumor growth. Moreover, YAP1(+) cells within SCLC tumors most closely transcriptionally resemble LCNEC rather than SCLC. This raises the possibility that the impact of YAP1 is not limited to transcriptional plasticity but histologic plasticity with a shift away from pure SCLC toward mixed histology neuroendocrine carcinomas less vulnerable to SCLC-directed therapies. While SCLC is increasingly appreciated as the destination for tumors undergoing transformation in response to therapeutic pressure, this is the first evidence, of which we are aware, that there is further histologic plasticity beyond SCLC, in this case to the equally recalcitrant YAP1-positive LCNEC state.

Several new classes of therapies, including antibody-drug conjugates and T-cell engagers, have altered the perception of the invincibility of relapsed SCLC. Several of the most promising targets for these agents – DLL3 and SEZ6 – are absent from the YAP1(+) populations, which may explain the infrequency of complete, durable responses. Nevertheless, YAP1(+) cells are not without their unique vulnerabilities, with robust expression of cell surface targets including B7-H3, TROP2, and HER2. Thus, barring a better-than-expected bystander effect with therapies targeting neuroendocrine-associated cell surface antigens, the most efficacious approaches to relapsed SCLC may rely on combination strategies, including those already in development (e.g. DLL3 + B7-H3 therapies used in concert).

We propose that, not only is YAP1 measurably expressed in many *relapsed* SCLCs, but that its presence underlies chemoresistance in the relapsed setting, as well as for the incomplete responses that typify even the most effective novel therapies for relapsed SCLC. If not specifically prevented or eliminated, *YAP1*(+) cells may underlie a persistent tumor compartment capable of adopting multiple states - neuroendocrine or non-neuroendocrine, epithelial or mesenchymal, even SCLC or LCNEC – contingent upon the most suitable state to evade the current therapy.

## Methods

### Transcriptional sequencing datasets

RNAseq analysis datasets from normal tissues were obtained from GTEx Portal (http://www.gtexportal.org) and tumor samples were retrieved from published relapsed SCLC datasets, including SCLC^17, 36, 52^.

### Cell culture

Human SCLC cell lines were purchased from ATCC. Cell lines were grown in RPMI with 10% fetal bovine serum and antibiotics and cultured at 37°C in a humidified chamber with 5% CO2. All cell lines were frequently tested for Mycoplasma. Cell lines demonstrated a range of phenotypic characteristics, from floating aggregates to spindle-like or cobblestone attached, but these were consistent with ATCC or previous reports. H69 and H69CPR cells were cultured and cell pellets were analyzed by RPPA, as described previously^53^. The same cells were injected subcutaneously into mice and tumors were harvested and analyzed by RPPA. SCLC cell lines (H1876 and H2195) were treated with 1 μM cisplatin for nine days. Cells were harvested on days 0, 2, 5, and 9 and lysates were analyzed by western blots for ASCL1 (43666S; Cell Signaling), YAP1, NOTCH2 (D76A6; Cell Signaling), and Actin as a loading control.

### Patient consent and tissue collection

Patients diagnosed with SCLC or a combined high-grade neuroendocrine carcinoma, including SCLC, at the University of Texas M.D. Anderson Cancer Center were selected on the basis of disease irrespective of age, gender or other clinical criteria. These patients underwent informed consent to Institutional Review Board (IRB)-approved protocol 2020-0412, PA13-0589, or PA14-0276. Core needle biopsies, EBUS biopsies, or surgical resections were collected into RPMI media with antibiotics and transported quickly to the research lab for sample preparation and single-cell RNAseq.

### Histology and IHC

Independent pulmonary pathologists carefully reviewed H&E images from LCNEC, SCLC, and combined histology tumors to accurately assess morphology. Immunohistochemistry (IHC) was performed with a Bond Max automated staining system (Leica Microsystems Inc., Vista, CA) using a Clinical Laboratory Improvement Amendments (CLIA)-certified YAP1 antibody. CLIA certified YAP1 antibody and protocol used by the MDACC Translational Molecular Pathology Department for clinical samples. Nuclear expression YAP1 was quantified using a 4-value intensity score (0, none; 1, weak; 2, moderate; and 3, strong) and the percentage (0%–100%) of reactivity. A final expression score (H-score) was obtained by multiplying the intensity and reactivity extension values (range, 0–300), as described previously^53, 54^. Scoring of IHC expression was performed in malignant cells and specifically in the NEC cells of the mixed histology tumors. IHC data were examined by two experienced pathologists (AS, LSS).

### CyTOF liquid biopsy processing

Peripheral blood mononuclear cells were isolated with Ficoll-Paque Plus density gradient centrifugation at 1300 RPM for 40 minutes without brake at room temperature. Cell suspensions were washed with PBS and centrifuged at 300 x g for 7 min. Cell pellets were then resuspended in 1-2 mL of PBS, counted with an automated cell counter and viability was assessed using Trypan Blue exclusion. For assessing viability during cytometry time of flight (CyTOF), cell pellets were subsequently incubated in 0.5 μM Cell-ID cisplatin (Fluidigm #201064 in PBS) for 5 min at room temperature. Cisplatin reactivity was quenched with 10% FBS RPMI media and cell suspensions were centrifuged for 5 min at 300 x g followed by another wash with RPMI media. Cell pellets were resuspended in PBS to reach 0.5 - 1 × 10^6^ cells/mL aliquots per sample and fixed with 1.6% paraformaldehyde (PFA) for 10 min at room temperature. Samples were centrifuged and washed twice at 500 × g for 5 min at 4 °C to remove PFA with Cell Staining Media (CSM, 0.5% w/v BSA, 0.02% w/v NaN_3_ in low-Barium PBS). Finally, cell pellets were resuspended in CSM and stored at -80°C until CyTOF analysis.

### CyTOF Antibody Staining

Samples were incubated with surface antibody master mix for 30 min at room temperature. After washing with CSM, cells were permeabilized with methanol for 10 min on ice. Following two washes with CSM, samples were then incubated with the intracellular antibody master mix for 30 min at room temperature, including YAP1 (phosphor-Y375; Abcam ab62751). Samples were washed twice with CSM and resuspended in PBS containing 1:5000 191Ir/193Ir MaxPar Nucleic Acid Intercalator (Fluidigm) and 1.6% PFA to stain DNA and stored at 4 °C overnight. Prior to CyTOF analysis, cells were washed once with CSM, twice with filtered double-distilled water, and finally resuspended (∼10^6^ stained cells/mL) in filtered double-distilled water containing normalization beads (EQ Beads, Fluidigm). Apart from antibody metal isotopes, we also recorded event length, barcoding channels, normalization beads, DNA (191Ir and 193Ir), and dead cells (195Pt and 196Pt). Samples were analyzed at the MDACC NORTH Campus Flow Cytometry and Cellular Imaging Core Facility.

### CyTOF Data Processing

Following data normalization, data transformation was achieved by using the inverse hyberbolic sine (ArcSinh) function with a cofactor of 552. FCS files were uploaded and analyzed on Cytobank for initial data processing and traditional cytometry statistics. Beads, daoublets, dead (cisplatin-positive) and apoptotic (cleaved PARP) cells were removed by gating and CD45-CD56+ cells were gated for all subsequent single-cell analysis.

### Patient Methylation Analyses

Using publicly available reduced representation bisulfite sequencing (RRBS) of SCLC patient formalin fixed paraffin embedded (FFPE) tissue or circulating tumor (ct)DNA^52^, YAP1 promoter methylation was evaluated. Additionally, both RRBS and RNAseq were performed on the FFPE patient biopsies.

### Single-cell RNAseq Tissue Dissociation and Sequencing

Tissues were digested gently overnight at 37°C using collagenase A (1 mg/ml). Red blood cell lysis was performed, as needed. Cells were counted and only samples with >50% viability were submitted for single-cell RNA sequencing at the MD Anderson’s CPRIT SINGLE Core facility.

### Single-cell RNAseq Analyses

Raw data for single cell datasets were processed using the Cell Ranger v3.1.0^55^ to obtain the unique molecular identifier (UMI) data matrix. Cells with less than 300 detectable genes were filtered out. Samples from different batches were normalized and integrated following the sample integration pipeline in SEURAT v3^56, 57^. The integrated datasets were processed using SEURAT, including selecting significant PCA components for dimension reduction, Uniform Manifold Approximation and Projection (UMAP) conversion and visualization, and density-based clustering for subpopulation discovery. Cells with high UMI count were excluded as potential doublets. The cell subpopulations were identified and annotated using SingleR package^58^ with manual curation. The gene expression levels were visualized on UMAP by either colored feature plot or binary plot. For each gene, the expression status is defined as positive if the cell has non-zero expression value, or negative otherwise. Epithelial-mesenchymal transition (EMT) score was calculated based on the EMT signature as described previously^10, 20, 24^. Similarly, YAP/TAZ target score^18^, NE score^25^, Cellular (<u>Cyto</u>) <u>T</u>rajectory <u>R</u>econstruction <u>A</u>nalysis using gene <u>C</u>ounts and <u>E</u>xpression (CytoTRACE) score, and SenMayo^35^ scores were all calculated as described previously. The cancer subtypes were classified with a custom model built by CellTypist^32^, which was trained on selected patient derived xenograft (PDX) models with known subtypes. The inferred copy number status was identified by the CopyKAT package.

### Pseudotime Analysis

Pseudotime trajectory analysis was performed using the Monocle2 R package (version 2.22.0) from the expression count matrix^59^. A CellDataSet (CDS) object was constructed using the newCellDataSet() function with the negative binomial distribution specified to model count-based data. Size factors and dispersion estimates were computed using estimateSizeFactors() and estimateDispersions(), respectively, to normalize for sequencing depth and account for gene-level variability. Genes were based on dispersion statistics obtained via dispersionTable() (e.g., mean_expression > 0.1 and empirical_dispersion > fit_dispersion)^60^.

Dimensionality reduction was carried out using the DDRTree method via reduceDimension(method = “DDRTree”), followed by cell ordering in pseudotime using orderCells(). To define the root state of the trajectory, we selected a biologically relevant root cell or state based on known marker gene expression patterns. Specifically, we visualized candidate genes using plot_cell_trajectory(), and assigned the root state via the root_state parameter in orderCells()^61^. Gene expression dynamics across pseudotime were visualized using plot_genes_in_pseudotime() and plot_pseudotime_heatmap() to identify key regulators of cell state transitions.

### Statistical Analyses

All statistic and bioinformatics analyses were performed using R. Paired and un-paired two-sample t-tests were used for two group comparisons on paired and unpaired experimental design experiments. Non-parametric Wicoxon rank-sum test (Mann-Whitney U test) was used for bulk RNAseq analysis of naïve vs. relapsed samples. Pearson and Spearman correlations were used for correlating genomic and proteomic measurements, as well as correlating drug-screening data. In all the analyses, p<0.05 was considered statistically significant.

### Data Set Availability

Publicly available data were obtained from relapsed SCLC patient tumors^16, 17, 36^. Patient single-cell RNA-seq data generated in this study have been deposited into the NCBI’s GEO with accession number GSE269942.

## Acknowledgements

We thank the patients who participated in this study, as well as their families. This work was supported by: The NIH/NCI CCSG P30-CA016672 (MD Anderson Flow Cytometry and Cellular Imaging Facility, the MD Anderson Bioinformatics Shared Resource, Advanced Technology Genome Core (ATGC), and the MD Anderson Institutional Tissue bank (ITB); This research was performed at the Single Cell Genomics Core Facility, which is supported in part by CPRIT Single Core grant RP180684; NIH 1S10OD024977-01 award to the ATGC for the NovaSeq6000; NIH/NCI T32-CA009666 (B. Zhang); The University of Texas-Southwestern and MD Anderson Cancer Center Lung SPORE P50-CA070907 (J. Wang, J.V. Heymach, C.M. Gay, L.A. Byers); NIH/CHI R01-CA299261 (C.M. Gay), NIH/NCI R01-CA207295 (L.A. Byers); NIH/NCI U01-CA213273 (J.V. Heymach, L.A. Byers); NIH/NCI R50-CA243698 (C.A. Stewart); NIH/NCI U01-CA256780 (J.V. Heymach, L.A. Byers); NIH/NCI U24-CA213274 (L.A. Byers); The Department of Defense LC210510 (L.A. Byers); the American Lung Association Pierre Massion Lung Cancer Discovery Award (L.A.B.), the LUNGevity Foundation 2020-02 (CM. Gay); CPRIT RP210159 (C.M. Gay); NETRF (C.M. Gay); Through generous philanthropic contributions to The University of Texas MD Anderson Lung Cancer Moon Shot Program (J.V. Heymach, L.A. Byers, C.M. Gay); and Rexanna’s Foundation for Fighting Lung Cancer (J.V. Heymach, L.A. Byers, C.M. Gay). This study was supported by the Translational Molecular Pathology-Immunoprofiling Lab (TMP-IL) Moon Shots Platform at the Department Translational Molecular Pathology, the University of Texas MD Anderson Cancer Center with special thanks to Wei Lu, Mei Jiang, Khaja Khan and Jianling Zhou for their technical assistance in the completion of this project. We would also especially like to thank A.R.K., L.W.Y., M.J.A., R.B.N., K.E.N., J.O., J.K.R., B.& B.N, C.K., P.C.B., S.S., S.R. and W.A.B. for their philanthropic support of these projects.

## Competing Interests

C.M.G. has speaking engagements with ACHL, AstraZeneca, BeiGene, Dava Oncology, IDEOlogy, IDR, MJH, OncLive, PeerView, PER, Targeted Healthcare, UpToDate; serves on the Advisory Board/Steering Committee for Abdera, Amgen, AstraZeneca, BMS, Boehringer Ingelheim, Daiichi Sankyo, G1, Jazz, Monte Rosa, Prophet, Roche/Genentech; consults for Catalyst and Kisoji. L.A.B. serves on consulting/advisory boards for Merck Sharp & Dohme Corp., Arrowhead Pharmaceuticals, Chugai Pharmaceutical Co., AstraZeneca Pharmaceuticals, Genetech Inc., BeiGene, AbbVie, Jazz Pharmaceuticals, Puma Biotechnology, Amgen, Daiichi Sankyo, Novartis and receives research funding from AstraZeneca Pharmacueticals, Amgen, and Circle Pharma. Otherwise, there are no pertinent financial or non-financial conflicts of interest to report.

## Author Contributions

C.A.S., C.M.G. conceived the project, analyzed and interpreted the data, and wrote the manuscript; C.A.S., R.W., A.T., B.Z., Y.Y., and M.B. performed experiments and interpreted results; C.A.S., K.R., B.L.R., A.G.S. interpreted results; Y.X., L.D., P.D.R., L.S.S., L.K., J.G., L.K., J.W. contributed to the analysis and interpretation of data; C.M.G., B.Z., R.W., A.H., W.L., L.K., J.V.H., L.A.B. contributed to the acquisition of data, administrative, and/or material support. All authors contributed to the writing, review, and/or revision of the manuscript.

## References

1. Liu SV, Reck M, Mansfield AS, et al. Updated Overall Survival and PD-L1 Subgroup Analysis of Patients With Extensive-Stage Small-Cell Lung Cancer Treated With Atezolizumab, Carboplatin, and Etoposide (IMpower133). J Clin Oncol 2021;39:619–630.

2. Paz-Ares L, Borghaei H, Liu SV, et al. Efficacy and safety of first-line maintenance therapy with lurbinectedin plus atezolizumab in extensive-stage small-cell lung cancer (IMforte): a randomised, multicentre, open-label, phase 3 trial. Lancet 2025;405:2129–2143.

3. Paz-Ares L, Chen Y, Reinmuth N, et al. Durvalumab, with or without tremelimumab, plus platinum-etoposide in first-line treatment of extensive-stage small-cell lung cancer: 3-year overall survival update from CASPIAN. ESMO Open 2022;7:100408.

4. Gay CM, Stewart CA, Park EM, et al. Patterns of transcription factor programs and immune pathway activation define four major subtypes of SCLC with distinct therapeutic vulnerabilities. Cancer Cell 2021;39:346–360 e347.

5. Xie M, Chugh P, Broadhurst H, et al. Durvalumab (D) plus platinum-etoposide (EP) in 1L extensive-stage small-cell lung cancer (ES-SCLC): Exploratory analysis of SCLC molecular subtypes in CASPIAN. ONCOLOGY RESEARCH AND TREATMENT. KARGER ALLSCHWILERSTRASSE 10, CH-4009 BASEL, SWITZERLAND; 2022;45:145–145.

6. Liu S, Mok T, Mansfield A, et al. VP5-2021: IMpower133: gene expression (GE) analysis in long-term survivors (LTS) with ES-SCLC treated with first-line carboplatin and etoposide (CE)±atezolizumab (atezo). Annals of Oncology 2021;32:1063–1065.

7. McColl K, Wildey G, Sakre N, et al. Reciprocal expression of INSM1 and YAP1 defines subgroups in small cell lung cancer. Oncotarget 2017;8:73745–73756.

8. Zanconato F, Cordenonsi M, Piccolo S. YAP/TAZ at the Roots of Cancer. Cancer Cell 2016;29:783–803.

9. Rudin CM, Poirier JT, Byers LA, et al. Molecular subtypes of small cell lung cancer: a synthesis of human and mouse model data. Nat Rev Cancer 2019;19:289–297.

10. Stewart CA, Diao L, Xi Y, et al. YAP1 Status Defines Two Intrinsic Subtypes of LCNEC with Distinct Molecular Features and Therapeutic Vulnerabilities. Clinical cancer research : an official journal of the American Association for Cancer Research 2024;30:4743–4754.

11. Owonikoko TK, Dwivedi B, Chen Z, et al. YAP1 Expression in SCLC Defines a Distinct Subtype With T-cell-Inflamed Phenotype. J Thorac Oncol 2021;16:464–476.

12. Ng J, Cai L, Girard L, et al. Molecular and Pathologic Characterization of YAP1-Expressing Small Cell Lung Cancer Cell Lines Leads to Reclassification as SMARCA4-Deficient Malignancies. Clinical cancer research : an official journal of the American Association for Cancer Research 2024;30:1846–1858.

13. Baine MK, Hsieh MS, Lai WV, et al. SCLC Subtypes Defined by ASCL1, NEUROD1, POU2F3, and YAP1: A Comprehensive Immunohistochemical and Histopathologic Characterization. J Thorac Oncol 2020;15:1823–1835.

14. Wu Q, Guo J, Liu Y, et al. YAP drives fate conversion and chemoresistance of small cell lung cancer. Sci Adv 2021;7:eabg1850.

15. Ireland AS, Micinski AM, Kastner DW, et al. MYC Drives Temporal Evolution of Small Cell Lung Cancer Subtypes by Reprogramming Neuroendocrine Fate. Cancer Cell 2020;38:60–78 e12.

16. Jiang L, Huang J, Higgs BW, et al. Genomic Landscape Survey Identifies SRSF1 as a Key Oncodriver in Small Cell Lung Cancer. PLoS Genet 2016;12:e1005895.

17. George J, Maas L, Abedpour N, et al. Evolutionary trajectories of small cell lung cancer under therapy. Nature 2024;627:880–889.

18. Wang Y, Xu X, Maglic D, et al. Comprehensive Molecular Characterization of the Hippo Signaling Pathway in Cancer. Cell reports 2018;25:1304–1317 e1305.

19. Hodgkinson CL, Morrow CJ, Li Y, et al. Tumorigenicity and genetic profiling of circulating tumor cells in small-cell lung cancer. Nature medicine 2014;20:897–903.

20. Stewart CA, Gay CM, Xi Y, et al. Single-cell analyses reveal increased intratumoral heterogeneity after the onset of therapy resistance in small-cell lung cancer. Nat Cancer 2020;1:423–436.

21. Zhang B, Stewart CA, Wang Q, et al. Dynamic expression of Schlafen 11 (SLFN11) in circulating tumour cells as a liquid biomarker in small cell lung cancer. Br J Cancer 2022;127:569–576.

22. Muhammad JS, Guimei M, Jayakumar MN, et al. Estrogen-induced hypomethylation and overexpression of YAP1 facilitate breast cancer cell growth and survival. Neoplasia 2021;23:68–79.

23. Chen H, Deng C, Gao J, et al. Integrative spatial analysis reveals tumor heterogeneity and immune colony niche related to clinical outcomes in small cell lung cancer. Cancer Cell 2025.

24. Byers LA, Diao L, Wang J, et al. An epithelial-mesenchymal transition gene signature predicts resistance to EGFR and PI3K inhibitors and identifies Axl as a therapeutic target for overcoming EGFR inhibitor resistance. Clinical cancer research : an official journal of the American Association for Cancer Research 2013;19:279–290.

25. Zhang W, Girard L, Zhang YA, et al. Small cell lung cancer tumors and preclinical models display heterogeneity of neuroendocrine phenotypes. Transl Lung Cancer Res 2018;7:32–49.

26. Lin G, Gao Z, Wu S, et al. scRNA-seq revealed high stemness epithelial malignant cell clusters and prognostic models of lung adenocarcinoma. Sci Rep 2024;14:3709.

27. Groves SM, Ildefonso GV, McAtee CO, et al. Archetype tasks link intratumoral heterogeneity to plasticity and cancer hallmarks in small cell lung cancer. Cell Syst 2022;13:690–710 e617.

28. Groves SM, Quaranta V. Quantifying cancer cell plasticity with gene regulatory networks and single-cell dynamics. Front Netw Physiol 2023;3:1225736.

29. Ma W, Zhou T, Song M, et al. Genomic and transcriptomic profiling of combined small-cell lung cancer through microdissection: unveiling the transformational pathway of mixed subtype. J Transl Med 2024;22:189.

30. Li J, Hubisz MJ, Earlie EM, et al. Non-cell-autonomous cancer progression from chromosomal instability. Nature 2023;620:1080–1088.

31. Jiang Y, Xie J, Cheng Q, et al. Comprehensive genomic and spatial immune infiltration analysis of survival outliers in extensive-stage small cell lung cancer receiving first-line chemoimmunotherapy. Int Immunopharmacol 2024;141:112901.

32. Xu C, Prete M, Webb S, et al. Automatic cell-type harmonization and integration across Human Cell Atlas datasets. Cell 2023;186:5876–5891 e5820.

33. Dominguez Conde C, Xu C, Jarvis LB, et al. Cross-tissue immune cell analysis reveals tissue-specific features in humans. Science 2022;376:eabl5197.

34. Weiler P, Lange M, Klein M, et al. CellRank 2: unified fate mapping in multiview single-cell data. Nat Methods 2024;21:1196–1205.

35. Saul D, Kosinsky RL, Atkinson EJ, et al. A new gene set identifies senescent cells and predicts senescence-associated pathways across tissues. Nat Commun 2022;13:4827.

36. Wagner AH, Devarakonda S, Skidmore ZL, et al. Recurrent WNT pathway alterations are frequent in relapsed small cell lung cancer. Nat Commun 2018;9:3787.

37. Drapkin BJ, George J, Christensen CL, et al. Genomic and Functional Fidelity of Small Cell Lung Cancer Patient-Derived Xenografts. Cancer Discov 2018;8:600–615.

38. Grunblatt E, Wu N, Zhang H, et al. MYCN drives chemoresistance in small cell lung cancer while USP7 inhibition can restore chemosensitivity. Genes Dev 2020;34:1210–1226.

39. Hosea R, Hillary S, Naqvi S, et al. The two sides of chromosomal instability: drivers and brakes in cancer. Signal Transduct Target Ther 2024;9:75.

40. Ahn MJ, Cho BC, Felip E, et al. Tarlatamab for Patients with Previously Treated Small-Cell Lung Cancer. N Engl J Med 2023;389:2063–2075.

41. Mountzios G, Sun L, Cho BC, et al. Tarlatamab in Small-Cell Lung Cancer after Platinum-Based Chemotherapy. N Engl J Med 2025.

42. Chandana SR, Choudhury NJ, Dowlati A, et al. First-in-human study of ABBV-706, a seizure-related homolog protein 6 (SEZ6)–targeting antibody-drug conjugate (ADC), in patients (pts) with advanced solid tumors. American Society of Clinical Oncology; 2024.

43. Rudin CM, Ahn M-J, Johnson ML, et al. OA04.03 Ifinatamab Deruxtecan (I-DXd) in Extensive-Stage Small Cell Lung Cancer (ES-SCLC): Interim Analysis of Ideate-lung01. 2024 World Conference on Lung Cancer. San Diego, CA: 2024.

44. Wang J, Duan J, Wu L, et al. OA04.06 Efficacy and Safety of HS-20093 in Extensive Stage Small Cell Lung Cancer in a Multicenter, Phase 1 Study (ARTEMIS-001) 2024 World Conference on Lung Cancer. San Diego, CA: 2024.

45. Dowlati A, Chiang AC, Cervantes A, et al. OA04.04 Sacituzumab Govitecan as Second-Line Treatment in Patients with Extensive Stage Small Cell Lung Cancer 2024 World Conference on Lung Cancer. San Diego, CA: 2024.

46. Wang J, Wu L, Li X, et al. OA04.05 SHR-A1921, A TROP-2 Targeted Antibody-Drug Conjugate (ADC), In Patients (pts) with Advanced Small-Cell Lung Cancer (SCLC) 2024 World Conference on Lung Cancer. San Diego, CA: 2024.

47. Wermke M, Felip E, Kuboki Y, et al. First-in-human dose-escalation trial of BI 764532, a delta-like ligand 3 (DLL3)/CD3 IgG-like T-cell engager in patients (pts) with DLL3-positive (DLL3+) small-cell lung cancer (SCLC) and neuroendocrine carcinoma (NEC). American Society of Clinical Oncology; 2023.

48. Choudhury N, Jain P, Dowlati A, et al. 698P Interim results from a phase I/II study of HPN328, a tri-specific, half-life (T1/2) extended DLL3-targeting T cell engager in patients (pts) with small cell lung cancer (SCLC) and other neuroendocrine neoplasms (NEN). Annals of Oncology 2023;34:S486.

49. Mikami H, Feng S, Naoi S, et al. A DLL3/CD3/CD137 trispecific T cell engager shows potent antitumor activity in small cell lung cancer models. Cancer Research 2023;83:1872–1872.

50. Zhou D, Byers LA, Sable B, et al. Clinical Pharmacology Profile of AMG 119, the First Chimeric Antigen Receptor T (CAR-T) Cell Therapy Targeting Delta-Like Ligand 3 (DLL3), in Patients with Relapsed/Refractory Small Cell Lung Cancer (SCLC). J Clin Pharmacol 2024;64:362–370.

51. Legend Biotech Announces FDA Clearance of IND Application for LB2102 in Extensive Stage Small Cell Lung Cancer. 2022 Available at https://legendbiotech.com/legend-news/legend-biotech-announces-fda-clearance-of-ind-application-for-lb2102-in-extensive-stage-small-cell-lung-cancer/.

52. Heeke S, Gay CM, Estecio MR, et al. Tumor- and circulating-free DNA methylation identifies clinically relevant small cell lung cancer subtypes. Cancer Cell 2024;42:225–237 e225.

53. Byers LA, Wang J, Nilsson MB, et al. Proteomic profiling identifies dysregulated pathways in small cell lung cancer and novel therapeutic targets including PARP1. Cancer discovery 2012;2:798–811.

54. Fujimoto J, Kadara H, Garcia MM, et al. G-protein coupled receptor family C, group 5, member A (GPRC5A) expression is decreased in the adjacent field and normal bronchial epithelia of patients with chronic obstructive pulmonary disease and non-small-cell lung cancer. J Thorac Oncol 2012;7:1747–1754.

55. Zheng GX, Terry JM, Belgrader P, et al. Massively parallel digital transcriptional profiling of single cells. Nat Commun 2017;8:14049.

56. Hao Y, Hao S, Andersen-Nissen E, et al. Integrated analysis of multimodal single-cell data. Cell 2021;184:3573–3587 e3529.

57. Satija R, Farrell JA, Gennert D, et al. Spatial reconstruction of single-cell gene expression data. Nat Biotechnol 2015;33:495–502.

58. Aran D, Looney AP, Liu L, et al. Reference-based analysis of lung single-cell sequencing reveals a transitional profibrotic macrophage. Nat Immunol 2019;20:163–172.

59. Qiu X, Mao Q, Tang Y, et al. Reversed graph embedding resolves complex single-cell trajectories. Nat Methods 2017;14:979–982.

60. Trapnell C, Cacchiarelli D, Grimsby J, et al. The dynamics and regulators of cell fate decisions are revealed by pseudotemporal ordering of single cells. Nat Biotechnol 2014;32:381–386.

61. Mao Q, Wang L, Goodison S, et al. Dimensionality Reduction Via Graph Structure Learning. Proceedings of the 21th ACM SIGKDD International Conference on Knowledge Discovery and Data Mining 2015.

