## Supplemental Tables and Figures for "YAP1 defines an emergent, plastic population of relapsed small cell lung cancer"

**Supplementary Materials**

**Supplemental Table 1. Biopsy details for single-cell RNAseq**

| Biopsy (MDA-) | Patient sex | Diagnosis | Type of collection | Biopsy site | YAP1 status | Cancer cell # | Cluster # |
|---|---|---|---|---|---|---|---|
| SC416A | M | SCLC | Core needle | Lung/Mediastinal | High | 1,972 | 5 |
| SC439A | F | SCLC | Core needle | Head and neck | High | 1,689 | 4 |
| SC135A | M | SCLC | Core needle | Liver | High | 5,325 | 5 |
| SC233A | M | SCLC | Core needle | Liver | High | 1,236 | 7 |
| SC140A | M | SCLC | Core needle | Liver | High | 1,706 | 6 |
| SC262A | M | SCLC/LCNEC | Resection | Lung | High | 864 | 5 |
| SC437A | M | SCLC | Core needle | Retroperitoneal | High | 1,354 | 4 |
| SC162B | F | SCLC | Core needle | Lung | High | 6,888 | 6 |
| SC370A | M | SCLC | Core needle | Adrenal | Low | 3,282 | 5 |
| SC151B | M | SCLC | Core needle | Adrenal | Low | 1,750 | 6 |
| SC363A | F | SCLC | Resection | Brain | Low | 2,092 | 5 |
| SC396A | M | SCLC | Core needle | Lung | Low | 860 | 2 |
| SC279A | M | SCLC/LCNEC | Resection | Brain | Low | 359 | 3 |
| SC145A | F | SCLC | Core needle | Head and neck | Low | 5,448 | 6 |
| SC142A | M | SCLC | Core needle | Lymph node | Low | 981 | 5 |

**Supplemental Table 2. Concordance between diagnosis and relapsed histology and YAP1 expression.**

| Patient ID | Diagnosis histology | Relapsed histology | YAP1 in small cell histology (naïve) | YAP1 in small cell histology (relapsed) | YAP1 in other histology (relapsed) | % *YAP1* by scRNAseq (relapsed) |
|---|---|---|---|---|---|---|
| KN-SC-17 | SCLC | SCLC | NA | Positive | NA | NA |
| KN-SC-19 | SCLC | SCLC | NA | Positive | NA | NA |
| ICON-070 | SCLC | SCLC/LCNEC | NA | Positive | Positive | NA |
| SC135 | SCLC | SCLC | NA | Positive | NA | 36.60 |
| SC145 | SCLC | SCLC | NA | Negative | NA | 0 |
| SC140 | SCLC | SCLC | Negative | Negative | NA | 22.81 |
| SC142 | SCLC | SCLC | NA | Negative | NA | 0 |
| SC151 | SCLC | SCLC | NA | Negative | NA | 0 |
| KN-SC-18 | SCLC | SCLC | NA | Negative | NA | NA |
| SC133 | SCLC | SCLC | NA | Negative | NA | NA |
| SC137 | SCLC | SCLC | NA | Negative | NA | NA |
| SC148 | SCLC | SCLC | NA | Negative | NA | NA |
| SC171 | SCLC | SCLC | NA | Negative | NA | NA |
| KN-SC-25 | SCLC | SCLC | NA | Negative | NA | NA |
| ICON-068 | SCLC | SCLC | NA | Negative | NA | NA |
| SC83 | SCLC | SCLC/LCNEC | NA | Negative | Positive | NA |
| SC155 | SCLC | SCLC/LUAD | NA | Negative | Positive | NA |
| ICON-136 | SCLC | SCLC/LCNEC | NA | Negative | Positive | NA |
| SC439 | SCLC | SCLC | Negative | NA | NA | 11.37 |
| SC233 | SCLC | SCLC | NA | NA | NA | 31.53 |

**Supplemental Table 3. Single-cell RNAseq percent expression of genes in relapsed SCLC biopsies.**

| Biopsy (MDA-) | Diagnosis | Subtype | *ASCL1* | *NEUROD1* | *POU2F3* | *YAP1* | *MYC* | *UCHL1* |
|---|---|---|---|---|---|---|---|---|
| SC416A | SCLC | A | 9.23% | 0 | 0 | 59.03% | 41.12% | 38.17% |
| SC439A | SCLC | A | 22.97% | 0 | 0 | 39.85% | 23.21% | 45.71% |
| SC135A | SCLC | P | 0.21% | 0 | 11.33% | 36.60% | 70.24% | 23.17% |
| SC233A | SCLC | A | 36.22% | 0 | 0 | 31.53% | 48.85% | 22.12% |
| SC140A | SCLC | A | 29.49% | 0 | 0.27% | 22.81% | 25.75% | 48.02% |
| SC262A | SCLC/LCNEC | A | 35.38% | 0 | 0 | 16.02% | 41.00% | 46.93% |
| SC437A | SCLC | A | 77.40% | 0 | 0 | 11.40% | 42.79% | 79.46% |
| SC162B | SCLC | N | 15.41% | 72.82% | 0 | 8.50% | 10.51% | 66.61% |
| SC370A | SCLC | N | 79.30% | 51.78% | 0 | 0 | 0 | 90.20% |
| SC151B | SCLC | A | 50.60% | 0 | 0 | 0 | 11.58% | 41.41% |
| SC363A | SCLC | I | 0 | 0 | 0 | 0 | 0 | 44.04% |
| SC396A | SCLC | I | 0 | 0 | 0 | 0 | 16.12% | 0 |
| SC279A | SCLC/LCNEC | A | 71.43% | 27.73% | 0 | 0 | 48.74% | 77.31% |
| SC145A | SCLC | N | 0 | 76.72% | 0 | 0 | 7.36% | 77.29% |
| SC142A | SCLC | I | 0 | 0.30% | 0 | 0 | 15.22% | 1.98% |

### Supplementary Figures

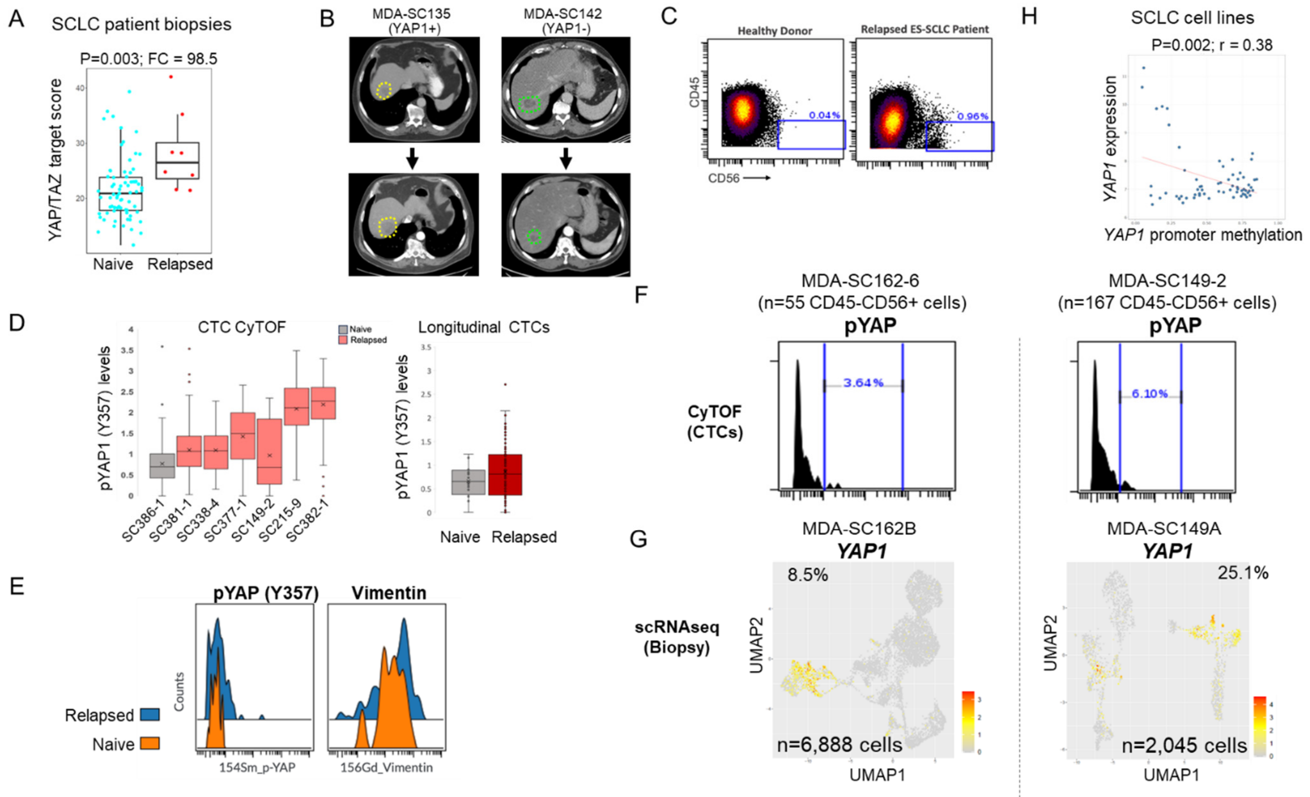


**Supplemental Fig. 1: YAP1 activity is elevated in relapsed SCLC and CTC protein levels are similar to paired patient biopsies.** A, YAP/TAZ target score in treatment naïve and relapsed biopsies. B, Patient scans demonstrating that a YAP1(+) tumor does not shrink following therapy (left), but a YAP1(-) tumor does (right). Dotted outline indicates tumor region. C, CyTOF analysis of liquid biopsies from a healthy donor and SCLC patient. A population of cells that are CD45-negative and CD56-positive are CTCs. D, CyTOF analysis of SCLC CTCs (25-254 cells per patient) demonstrate phospho (p)YAP1 (Y357) levels in relapsed patients (left). CyTOF analysis of YAP1 levels in longitudinal samples from the same patient at diagnosis and following relapse (right). E, pYAP1 and Vimentin are both higher in relapsed SCLC. F, CyTOF analysis of pYAP1 on CD45-CD56+ CTCs detected in liquid biopsies. G, scRNAseq analysis of YAP1 by scRNAseq in paired biopsies from the same patients reveal similar expression patterns. H, *YAP1* expression is negatively correlated with promoter methylation in SCLC cell lines.

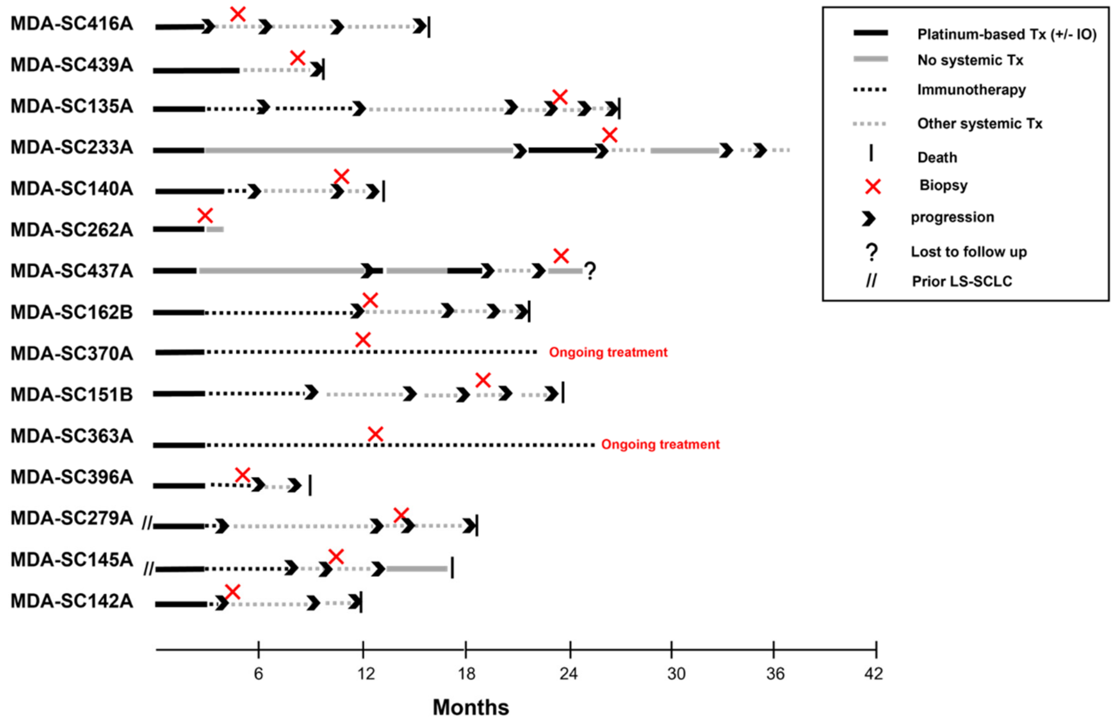


**Supplemental Fig. 2: Patient treatment course surrounding the biopsy analyzed by single-cell RNAseq.**

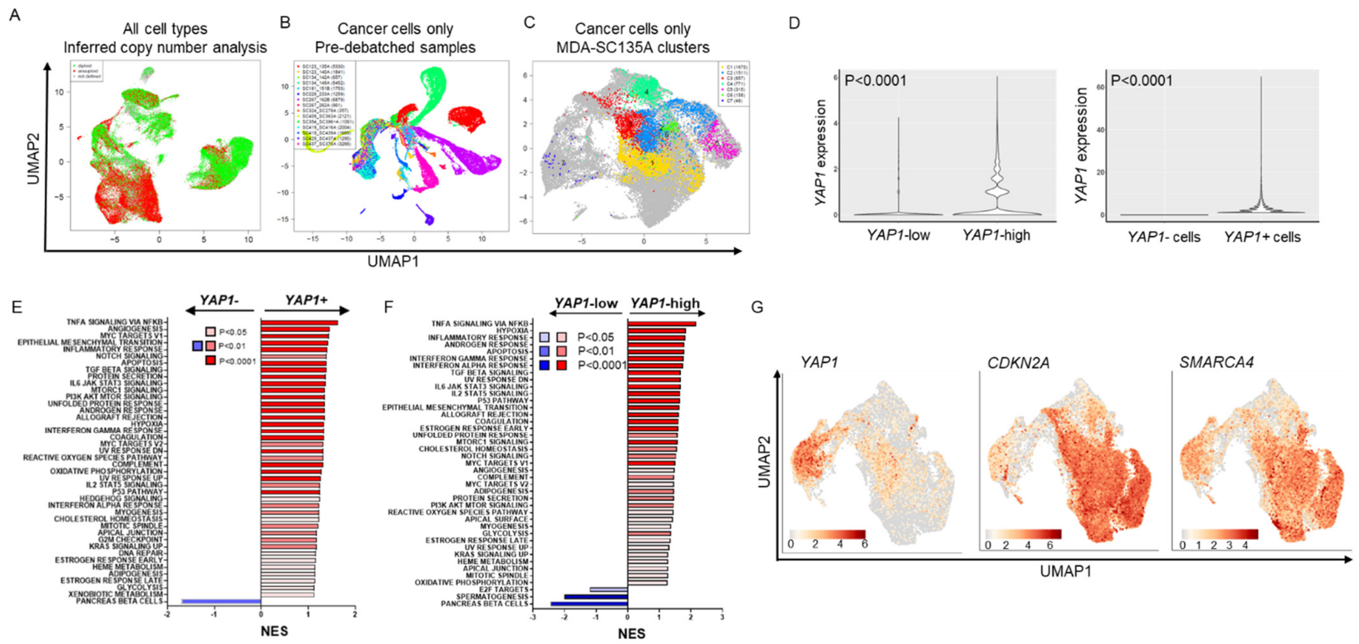

**Supplemental Fig. 3: Unique features of *YAP1*-high SCLC**. A, Inferred copy number analysis of all cells in relapsed SCLC confirms identification of cancer cell population clusters. B, Pre-debatched UMAP plot of cancer cells color-coded by patient demonstrates intertumoral heterogeneity. C, UMAP plot indicating clusters of cells from an individual biopsy cover numerous clusters in the pooled sample plot. D, *YAP1* expression in *YAP1*-high biopsies (left) and *YAP1*(+) cells (right). E, F, GSEA analysis of pathways enriched in YAP1(+) (E) or YAP1-high biopsies (F). G, UMAP feature plots demonstrating expression of *CDKN2A* and *SMARCA4* does not overlap with *YAP1* in relapsed SCLC biopsies.

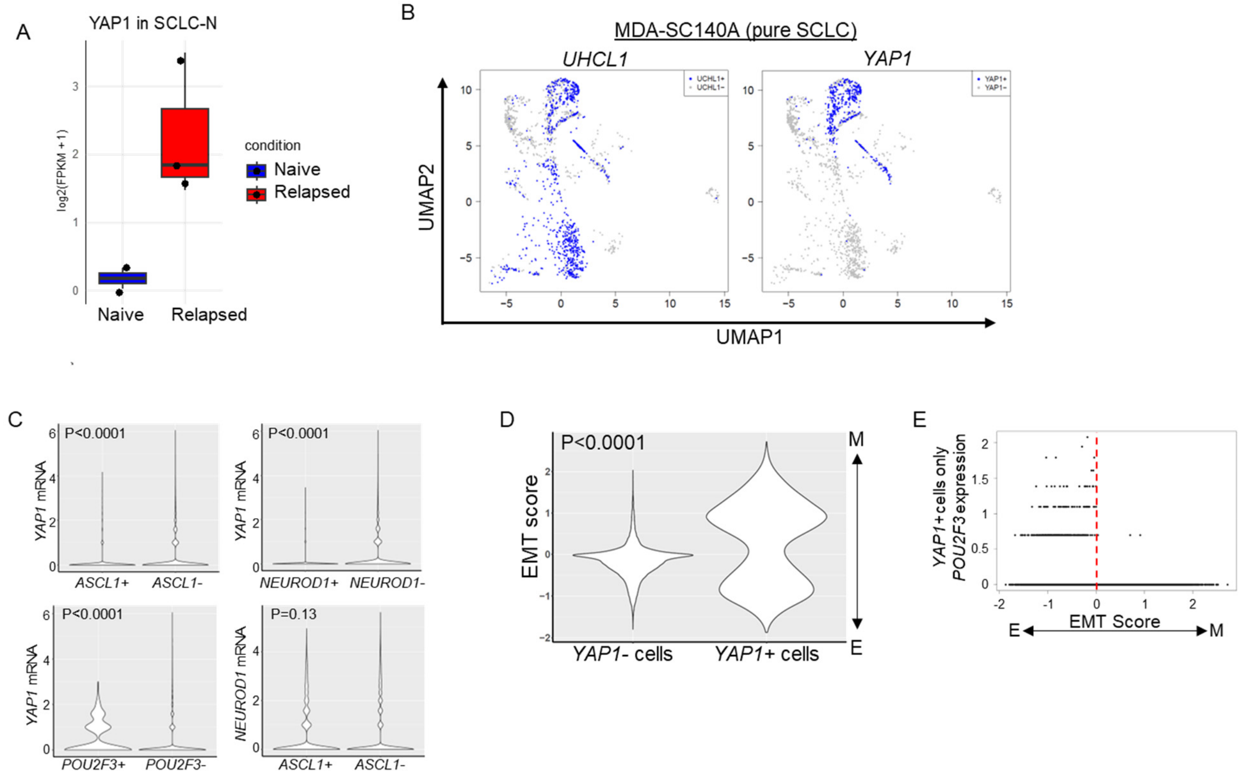


**Supplemental Fig. 4: Features of YAP1-high relapsed SCLC**. A, Bulk RNAseq analysis of *YAP1* in SCLC-N subtype patient tumors at treatment naïve and relapsed timepoints. B, Expression of *UHCL1* (NE gene) and *YAP1* in the same cell population in MDA-SC140A biopsy. C, Violin plots demonstrating co-expression of *YAP1* or *NEUROD1* in *ASCL1*, *NEUROD1*, or *POU2F3* positive and negative populations. D, EMT score range in *YAP1*(+) and (-) cells. E, *POU2F3* expression in *YAP1*(+) cells is associated with a low EMT score.

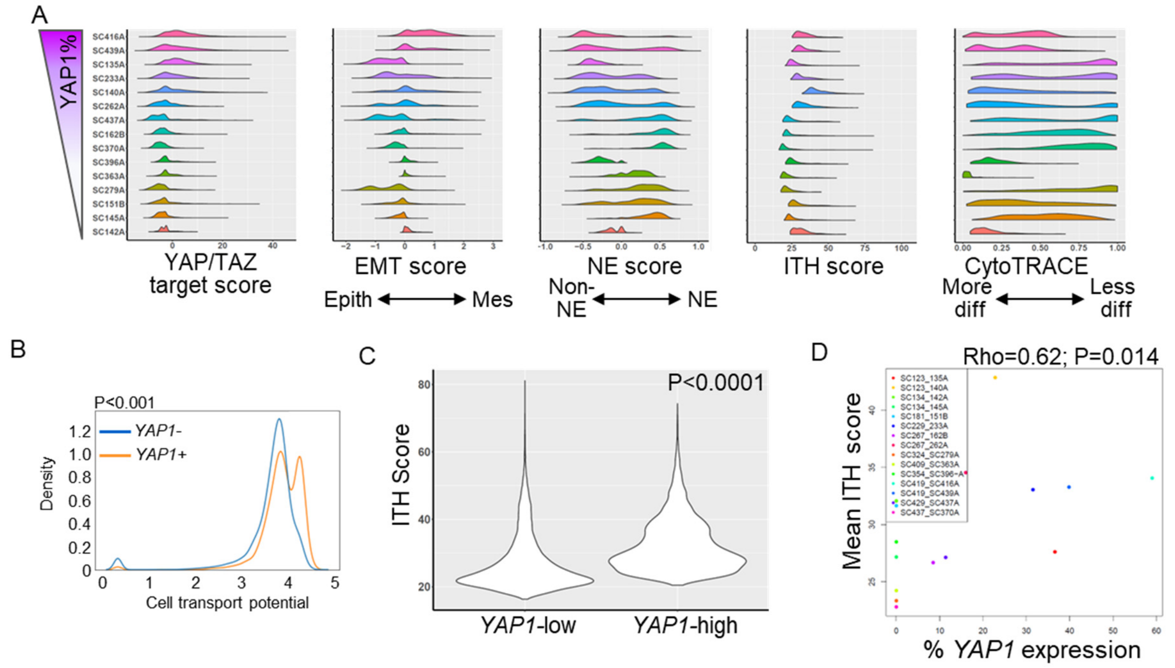


**Supplemental Fig. 5: YAP1 is associated with ITH and plasticity in relapsed SCLC.** A, Per sample distribution of common scores. Samples are ranked by *YAP1*%. B, Comparison of density and cell transport potential between *YAP1*(+) and (-) cells. C, Violin plot demonstrating increased ITH score in *YAP1*-high biopsies. D, Mean ITH score is directly correlated with % *YAP1* expression in relapsed biopsies (B, right).

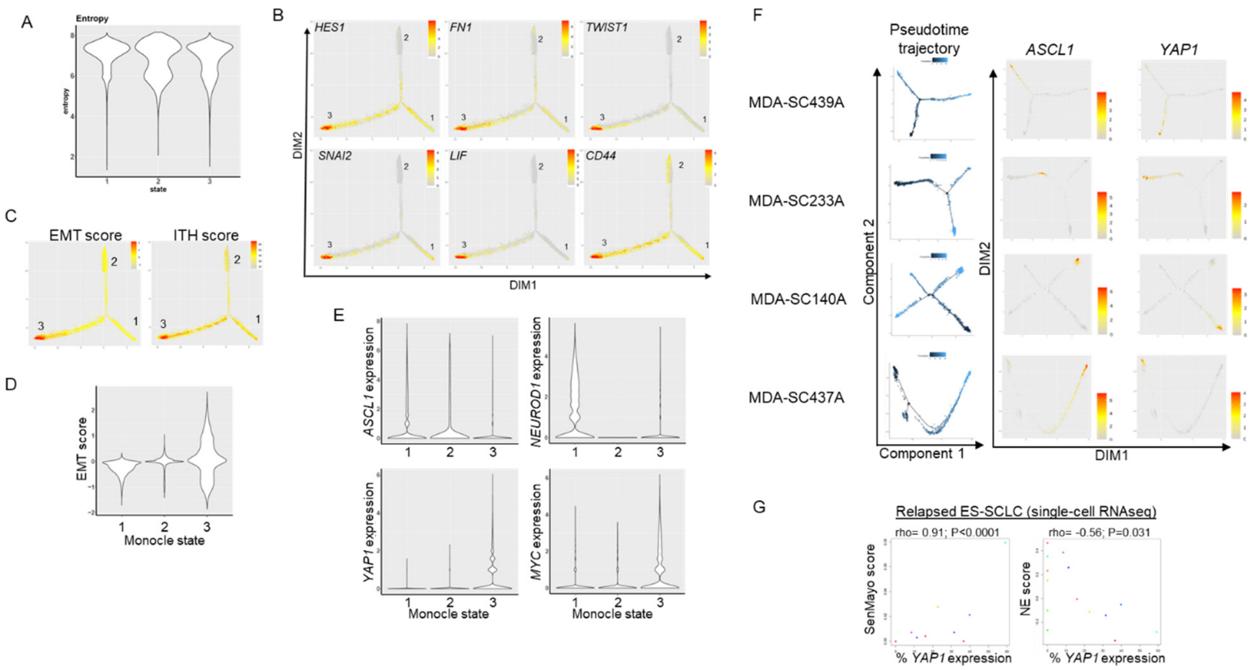


**Supplemental Fig. 6: Monocle state 3 is associated with ITH and EMT score in pooled patient biopsies**. A, Violin plot demonstrating transcriptional entropy across monocle states. B,C, DIM plots demonstrating expression patterns of plasticity genes (B) and EMT and ITH scores (C). D, Violin plots showing EMT score in monocle states. E, Violin plots showing expression of *ASCL1*, *NEUROD1*, *YAP1*, and *MYC* in monocle states. F, Pseudotime trajectory and DIM plots for individual patient biopsies demonstrating *ASCL1* and *YAP1* expression. G, Correlation between YAP1% expression and SenMayo or NE score.

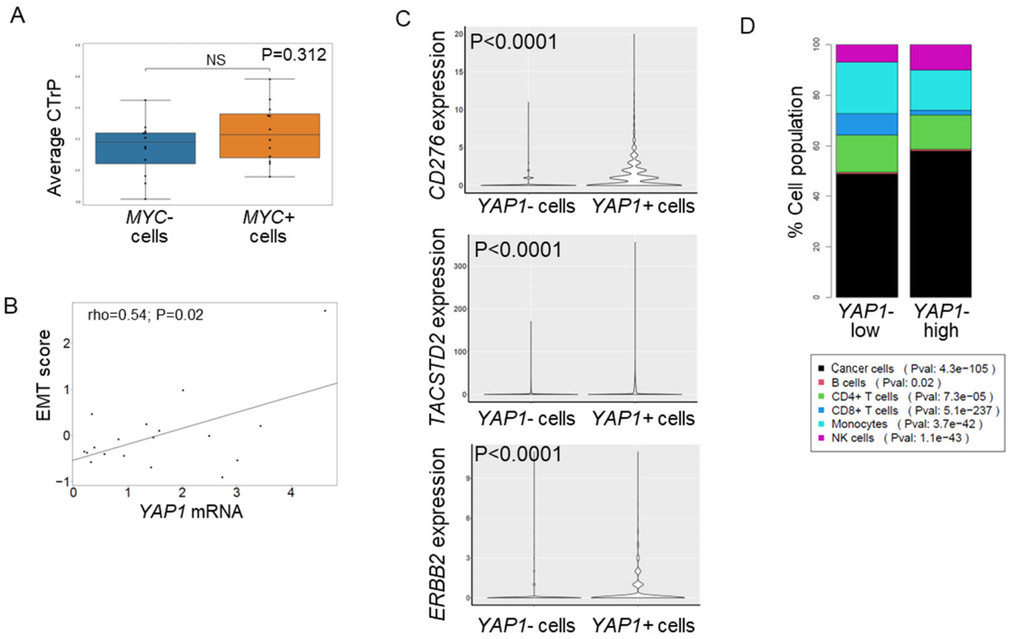


**Supplemental Fig. 7: YAP1 is co-expressed with surface targets and associated with NK cell abundance.** A, Average CTrP of *MYC*(+) and (-) populations was not different. B, *YAP1* and EMT score are positively correlated in relapsed patient biopsies by bulk RNAseq. C, Co-expression of *CD276*, *TACSTD2*, and *ERBB2* in *YAP1* (+) and (-) cells. D, *YAP1*-high biopsies have more cancer and NK cells, but less CD8+ T-cells.
